# Mapping Structure and Biological Functions within Mesenchymal Bodies using Microfluidics

**DOI:** 10.1101/514158

**Authors:** Sébastien Sart, Raphaël F.-X. Tomasi, Antoine Barizien, Gabriel Amselem, Ana Cumano, Charles N. Baroud

## Abstract

Organoids that recapitulate the functional hallmarks of anatomic structures comprise cell populations able to self-organize cohesively in 3D. However, the rules underlying organoid formation *in vitro* remain poorly understood because a correlative analysis of individual cell fate and spatial organization has been challenging. Here, we use a novel microfluidics platform to investigate the mechanisms determining the formation of organoids by human mesenchymal stromal cells that recapitulate the early steps of condensation initiating bone repair *in vivo*. We find that heterogeneous mesenchymal stromal cells self-organize in 3D in a developmentally hierarchical manner. We demonstrate a link between structural organization and local regulation of specific molecular signaling pathways such as NF-κB and actin polymerization, which modulate osteo-endocrine functions. This study emphasizes the importance of resolving spatial heterogeneities within cellular aggregates to link organization and functional properties, enabling a better understanding of the mechanisms controlling organoid formation, relevant to organogenesis and tissue repair.

## 1. Introduction

In recent years, organoids have emerged as powerful tools for basic research, drug screening and tissue engineering. The organoids formed *in vitro* show many features of the structural organization and the functional hallmarks of adult or embryonic anatomical structures^1^. In addition, the formation of organoids alleviates the need to perform animal studies and provide an attractive platform for robust quantitative studies on the mechanisms regulating organ homeostasis and tissue repair *in vivo*^1^. The formation of organoids usually starts with populations of stem cells. They are therefore expected to be heterogeneous since pluripotent stem cells (iPSCs or embryonic stem cells) have been shown to dynamically and stochastically fluctuate from ground to differentiated state^2,3^. In the same vein, LGR5^+^ intestinal stem cells are reported to contain several distinct populations^4^. As such the formation of organoids involves the inherent capacity of these heterogeneous populations to self-sorting and self-patterning in order to form an organized 3D architecture^5,6^. However, the rules underlying organoid formation as well as the contribution of intrinsic population heterogeneity to the organoid self-assembly remain poorly understood^5,6^. Consequently, there is a need for novel quantitative approaches at the single cell level to reliably understand the mechanisms of spatial tissue patterning in 3D organoids, for which microfluidic and quantitative image analysis methods are well suited.

In this work we use mesenchymal progenitors, alternatively named mesenchymal stromal cells (MSCs), which constitute a self-renewing population with the ability to differentiate into adipocytes, chondrocytes and osteoblasts^7^. Although human MSCs (hMSCs) express tight levels of undifferentiation markers (e.g., CD105, CD44, and Sca-1), they constitute a heterogeneous population of cells that exhibit considerable variation in their biophysical properties, epigenetic status, as well as the basal level of expression of genes related to differentiation, immune-regulation and angiogenesis^8,9,10^. Nonetheless their aggregation leads to the formation of highly cohesive 3D spherical structures (which we designate hereafter as mesenchymal bodies, MBs) with improved biological activities in comparison to 2D cultures^11^. However, little is known on how hMSCs self-organize or if the intrinsic heterogeneity of the population regulates MB formation and individual cell functions in 3D.

The self-aggregation of hMSCs into MBs can recapitulate the early stages of mesenchymal condensation and it promotes the secretion of paracrine molecules taking part in the process of ossification^12^. During mesenchymal condensation *in vivo*, mesenchymal progenitors self-aggregate and form dense cell-cell contacts that lead to the initiation of bone organogenesis through endochondral (necessitating a chondrogenic intermediate) and intramembranous (direct osteogenic differentiation) ossification^13^. In addition, the formation of these 3D mesenchymal bodies *in vivo* is associated with the secretion of important paracrine molecules such as prostaglandin E2 (PGE2) and vascular endothelial growth factor (VEGF), which participate to the recruitment of endogenous osteoblasts, osteoclasts and blood vessels leading to the initiation/restoration of bone homeostasis^14,15^. In these two ossification processes, the induction of NF-κB target genes such as cyclooxygenase 2 (COX-2), and their downstream products (e.g. PGE2 and VEGF), play a critical role of developmental regulators of ossification and bone healing^16,17,18^. However, while mesenchymal condensation is critical for bone organogenesis, there is still a limited understanding on how the cellular spatial organization within 3D mesenchymal bodies regulates the individual cells’ endocrine functions^19^.

In the present work, we interrogate the influence of phenotypic heterogeneity within a population of stem cells on the mechanisms of self-assembly and functional patterning within 3D organoids, using hMSCs as a model of heterogeneous progenitor cell population. This is performed using a novel microfluidic platform for high-density formation of mensenchymal bodies combined with the analysis of individual cells by quantitative image analysis. Our study reveals that the progenitor cell population self-assembles in a developmentally hierarchical manner, and that the structural cellular arrangement in mensenchymal bodies regulates the functional patterning in 3D, by modulating locally the activities of regulatory molecular signaling.

## 2. Results

### 2.1 Self-organization of hMSCs in 3D mesenchymal bodies

Human MSCs are known to constitute a heterogeneous population^8,9,10^. To examine the cellular diversity within the population, hMSCs were first characterized by their expression of membrane markers. Most of the hMSC population expresses CD73, CD90, CD105 and CD146, but not CD31 (Fig. 1.A-F). However, a deeper analysis of the flow cytometry data shows that the hMSC population contains cells of heterogeneous size (C.V. = 33-37%) (Fig. 1.G) and having a broad distribution in the expression of CD146 (Fig. 1.F). Of note, the CD146 level of expression was linked to the size of the cells: the highest levels of CD146 were found for the largest cells (Fig. 1.H). Similar correlations with cell size were also observed for CD73, CD90 and CD105 (Fig. S1). In addition, upon specific induction, the hMSC population used in this study successfully adopted an adipogenic (Fig. 1.I), or an osteogenic phenotype in 2D (Fig. 1.J), demonstrating their mesenchymal progenitor identity.

**Figure 1.**
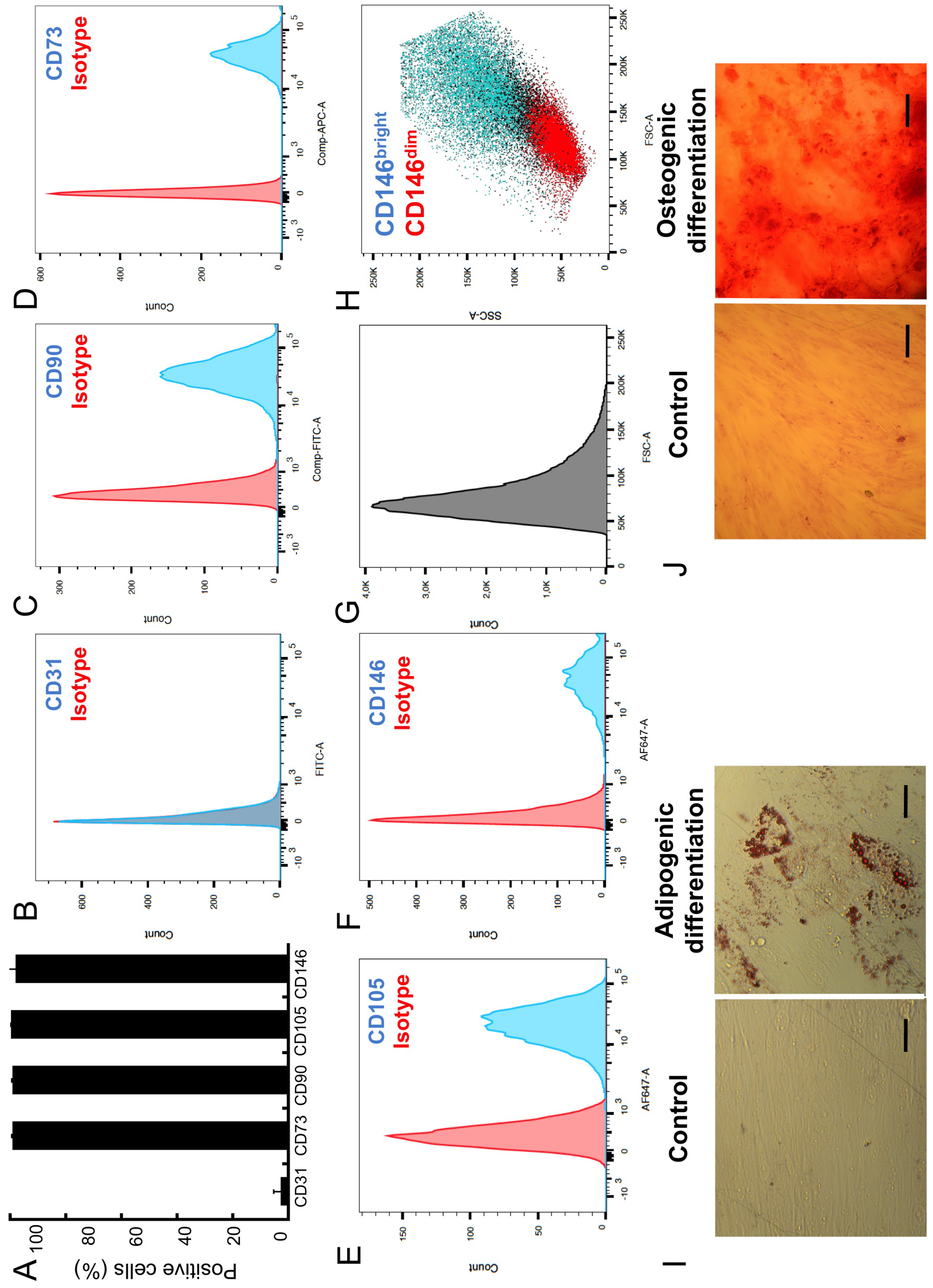
Characterization of the hMSC population. (A) Percentage of positive cells for CD 31, CD73, CD90, CD105 and CD146 (n=3). Representative histograms of the distribution of the CD31- (B), CD73- (C), CD90- (D), CD105- (E) and CD146- (F) level of expression. Representative histogram of the forward scatter distribution (G). Correlation between cell size (FSC and SSC) and the level of CD146 expression (H). (I) Representative images of hMSCs differentiated towards adipogenic lineage in 2D cultures (Oil red O staining). (J) Representative images of UC-hMSCs differentiated towards osteogenic lineage in 2D cultures (Alizarin red S staining). Scale bars are 50 μm.

To interrogate contribution of cellular heterogeneity in the self organization of hMSCs in 3D, mesenchymal bodies (MBs) were formed at high density on an integrated microfluidic chip. This was done by encapsulating cells into microfluidic droplets at a density of 200 cells per droplet. The drops were then immobilized in 250 capillary anchors^20^ in a culture chamber, as previously described (Fig. 2.A-B)^21^. The loading time for the microfluidic device was about 5 minutes, after which the typical time for complete formation of MBs was about 4 h, as obtained by measuring the time evolution of the projected area (Fig. 2.C-D) and circularity of individual MBs (Fig. 2.E). The protocol resulted in the formation of a single MB per anchor with an average diameter of 158 μm (Fig. 2.F). In addition, the complete protocol yielded the reproducible formation of a high density array of fully viable MBs ready for long-term culture (Fig. 2.G), as described previously^21^. Of interest, the coefficient of variation (C.V.) of the MB diameter distribution was significant lower than the C.V. of individual cell size (C.V. MB diameter = 13.3 % and C.V. cell size = 35 %), which demonstrates that despite its broad heterogeneity, the cell population was able to reproducibly self-organize within the MBs.

**Figure 2.**
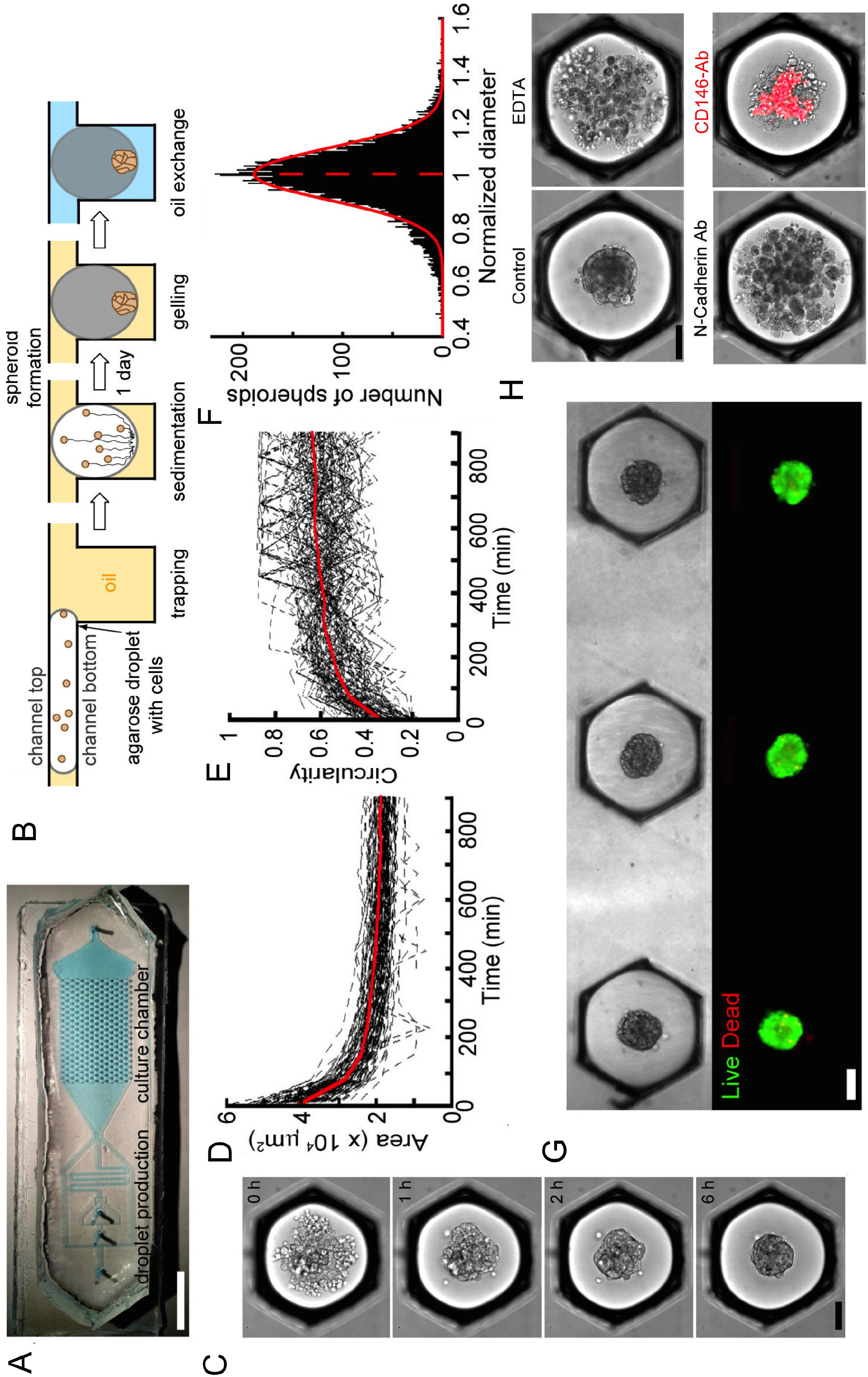
Formation of mesenchymal bodies on chip. (A) Chip design, scale bar is 1 cm. (B) Schematized side view of an anchor through the MB formation and culture protocol. (C) Representative time-lapse of a MB formation, scale bar is 100 μm. (D-E) Measurement of the time evolution of the projected area (D) and circularity of each aggregate (E), n_MBs_ = 120. (F) Distribution of the mesenchymal bodies’ diameter normalized by the mean of each chip (n = 10,072 MBs). (G) Representative images of MBs after agarose gelation and oil to medium phase change (top). The same MBs are stained with LIVE/DEAD^TM^ (bottom), scale bar is 100 μm. (H) Representative images of MBs formed in the presence of EDTA, a N-cadherin or a CD146-conjugated blocking antibody (the red color shows the position of the CD146 brightest cells, the dilution of the antibody was 1/100 and remain in the droplet for the whole experiment), scale bar is 100 μm. (I-J).

To gain insight on the cellular components required to initiate the self-organization of hMSCs in 3D, the MB formation was disrupted by altering cell-cell interactions. This was first performed by adding EDTA, a chelating agent of the calcium involved the formation of cadherin junctions, to the droplet contents. Doing so disrupted the MB formation, as shown in Fig. 2.H, where the projected area of the cells increased and the circularity decreased in the presence of EDTA compared to the controls. The role of N-cadherins among different types of cadherins was further specified by adding a blocking antibody in the droplets prior to MB formation. This also led to a disruption of the MB formation, demonstrating that N-cadherin homodimeric interactions are mandatory to initiate the process of hMSC aggregation. In the same vein, the addition of a CD146-conjugated blocking antibody also increased the projected area of the MBs (Fig. 2.H) as well reducing their circularity (Fig. 2.H), demonstrating that cell-cell interactions involving CD146 are also required during MB formation (Fig. 2.H). Of note, the brightest signal from the CD146 stained cells was located in the core of the cellular aggregates (Fig. 2.H).

While the population of hMSCs is constituted by cells of broad size and expressing different levels of undifferentiated markers (i.e. CD90, CD73, CD105 and CD146 are known to be down-regulated upon differentiation^22,23^), the cells are nonetheless capable to self-organize cohesively in 3D. To better understand how the heterogeneous cells organize within the MBs, we measured how the different cell types composing the population self-assembled spatially in 3D, by investigating the role of CD146. For this purpose, the CD146^dim^ and CD146^bright^ cells were separated from the whole hMSC population by flow cytometry (Fig. 3.A-B). The cells were then reseeded on chip for MB formation, after fluorescently labeling the brighter or the dimmer CD146 populations. Image analysis revealed that the CD146^bright^ cells were mostly located in the center of the cellular aggregates, while CD146^dim^ were found at the boundaries of the MBs (Fig. 3.C-D-E). As we found that the CD146^bright^ cells were significantly larger than the CD146^dim^ cells, the cells from the hMSC population were also separated based on their relative size (a parameter that also discriminates the CD90-, CD105- and CD73-^bright^ from the CD90-, CD105- and CD73-^dim^ cells, Fig. S1). After reseeding on chip, the MBs were composed of large cells in the core, while the smallest cells were located at the boundaries, as expected from the previous experiments (Fig. S2). It is well established that CD146^bright^ defines the most undifferentiated hMSCs^24,25,26^. The heterogeneity in level of commitment between the two subpopulations was therefore checked by RT-qPCR analysis, in order to quantify differences in the expression of differentiation markers. The analysis showed that the CD146^dim^ cells express higher level of osteogenic differentiation markers (i.e. RUNX-2) than the CD146^bright^ cells (Fig. 3. F-G).

**Figure 3.**
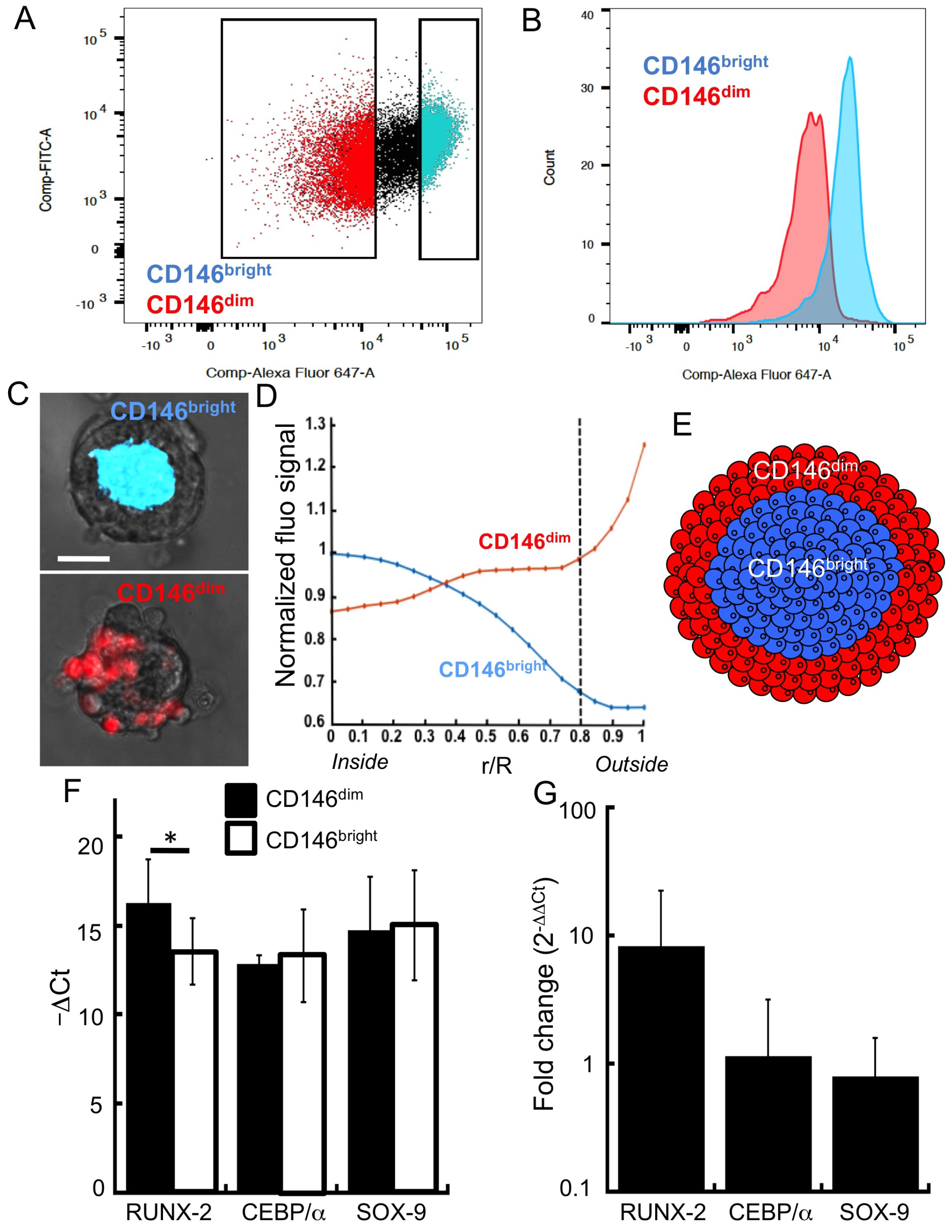
CD146 expression and cellular commitment regulate the structural organization of mesenchymal bodies. (A) Representative dot plot of the hMSC population separation based on the level of CD146: the CD146^bright^ constitute 20% of the population expressing the highest levels of CD146; the CD146^dim^ constitute the 20% of the population expressing the lowest levels of CD146. (B) Fluorescent signal distribution in the CD146^dim^ and CD146^birght^ populations after cell sorting. (C) After cell sorting the CD146^bright^ or the CD146^dim^ were stained with Vybrant^®^DIL, remixed together and allowed to form MBs. Representative images of the CD146^bright^ and CD146^dim^ within the MBs, scale bare is 50 μm. (D) The position of the CD146^bright^ and CD146^dim^ was quantified by correlating fluorescent signal of the different stained cells as a function of their radial position within the MBs. (E-F) RT-qPCR analysis of the relative RUNX-2, CEBP/α and SOX-9 expression to GADPH (ΔCt, D and relative RNA expression, E), in the CD146^bright^ and in the CD146^dim^ populations (n=3).

The level of RUNX-2 expression was also quantified at the protein level using immunocytochemistry and image analysis of the MBs on the microfluidic device, by developing a layer-by-layer description of the mesenchymal bodies. This mapping was constructed by estimating the boundaries of each cell in the image from a Voronoi diagram, built around the positions of the cell nuclei stained with DAPI (Fig. 4.A)^27,28^. These estimates were then used to associate the fluorescence signal from each cell with one of the concentric layers (Fig. 4.B). Such a mapping provides better resolution for discriminating the spatial heterogeneity of protein expression than simply assigning a fluorescent signal to a defined radial coordinate (Fig. S3). Moreover, the reliability of the measurements by quantitative image analysis was confirmed by performing several control experiments. In particular we verified (1) the specificity of the fluorescent labeling; (2) the absence of limitation for antibody diffusion; (3) the absence of the light path alteration in the 3D structure (Fig. S4). Consistent with the qPCR data we found that hMSCs located at the boundaries of the MBs expressed higher levels of the protein RUNX-2 than the cells located in the core (Fig. 4.C-D, Fig. S6 for individual experiments).

Thus, as CD146 defines the most undifferentiated and clonogeneic cells as well as regulating the tri-lineage differentiation potential of hMSCs^24,29^, the results indicate that hMSCs self-organize within MBs based on their initial commitment. The most undifferentiated and largest cells are found in the core (r/R < 0.8), while more differentiated cells positioned in the outer layers of the MBs (r/R > 0.8) (Fig. 4.C-D-E). In addition, these data reveal that hMSCs are conditioned *a-priori* to occupy a specific location within the mesenchymal bodies.

**Figure 4.**
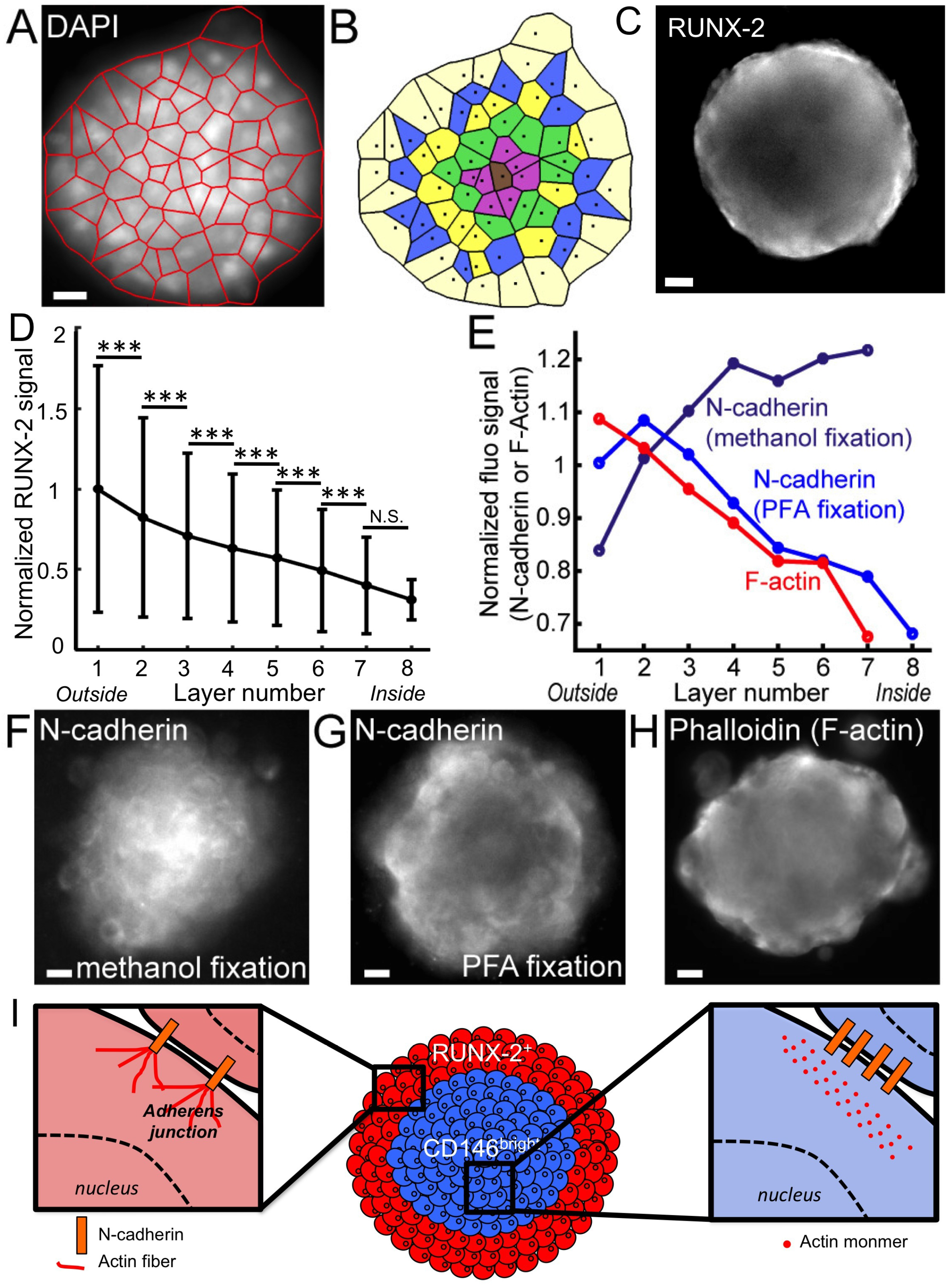
Quantitative mapping of RUNX-2 expression and structural organization in mesenchymal bodies. (A-B) The detection of nuclei within MBs enables the construction of a Voronoi diagram (A) that allows the identification of concentric cell layers (B) within the MBs. (C-D) Representative image (C) and quantitative analysis (D, error bars represent the standard deviation) of RUNX-2 staining within the cell layers of the MBs (N_chips_ = 3, n_MBs_ = 458). (E-H) Quantitative analysis (E) and representative images (F-H) of N-cadherin staining post methanol/acetone (F, N_chips_ = 3, n_MBs_ = 405), post PFA/Triton-X100 fixation and permeabilization (G, N_chips_ = 3, n_MBs_ = 649) and F-actin staining with phalloidin (H, N_chips_ = 3, n_MBs_ = 421). All scale bars are 20 μm. ***: p < 0.001; N.S.: non significant.

The commitment of hMSCs is known to regulate their level of expression and the type of cell-cell adhesion molecules^30,31^, which plays a fundamental role in the structural cohesion of the MBs (Fig. 2.H). For this reason we interrogated the organization of cell-cell junctions after MB formation, through measurements of the N-cadherin and F-actin fluorescent signal distribution. Two different protocols were used to discriminate several forms of N-cadherin interactions. First, PFA fixation and Triton-X100 permeabilization were used, since they were reported to retain in place only the detergent-insoluble forms of N-cadherin^32,33,34,35^. Alternatively, ice-cold methanol/acetone fixation and permabilization enabled the detection of all forms of N-cadherins^32,33,34,35^. The results show a higher density of total N-cadherins in the core of the MBs (Fig. 4.E-F), while a higher density of F-actin was found in the cell layers located near the edge of the MBs (Fig. 4.E,H). These results are consistent with the theories of cell sorting in spheroids that postulate that more adhesive cells (i.e. expressing more N-cadherin) should be located in the core, while more contractile cells (i.e. containing denser F-actin) are located at the edge of the MBs^36^. Moreover, our observations are in accordance with recent results demonstrating that hMSCs establishing higher N-cadherin interactions show reduced osteogenic commitment than hMSCs making fewer N-cadherin contacts, potentially through the modulation of Yap/Taz signaling and cell contractility^37^.

In contrast, the most triton-insoluble forms of N-cadherins were located at the boundaries of the hMSC aggregates (Fig. 4.E,G), at the same position as the cells containing the denser F-actin. These results demonstrate that different types of cellular interaction were formed between the core and the edges of the MBs, as a function of the degree of cell commitment that apparently stabilize the *adherens* junctions^38^ (Fig. 4.I).

### 2.2. Mapping the biological functions in mesenchymal bodies

We found above that hMSCs self organized in MBs as a function of their size and their degree of commitment, which may also regulate their endocrine functions^39^. Thus, we interrogated the functional consequences of the cellular organization in mesenchymal bodies, by investigating the distribution of VEGF and PGE2 producing cells.

The specific production of COX-2, VEGF and two other molecules regulating bone homeostasis such as tumor necrosis factor-inducible gene 6 (TSG-6)^40^ and stanniocalcin 1 (STC-1)^41^ was evaluated by RT-qPCR analysis. An increased transcription (20 to 60 fold) of these molecules was measured in 3D in comparison to the monolayer culture (Fig. 5.A-B). Consistent with this observation, while very limited level of secreted PGE-2 and VEGF was measured by ELISA in 2D culture, they were significantly increased (by about 15 fold) upon the aggregation of hMSCs in 3D (Fig. 5.C). In addition, to interrogate the specific role of COX-2 (the only inducible enzyme catalyzing the conversion of arachidonic acid into prostanoids^42^) in PGE-2 and VEGF production, indomethacin (a pan-COX inhibitor) was added to the culture medium. Indomethacin abrogated the production of PGE-2 and it significantly decreased VEGF secretion (Fig. 5.C), which suggests an intricate link between COX-2 expression and the secretion of these two molecules^43,44^.

**Figure 5.**
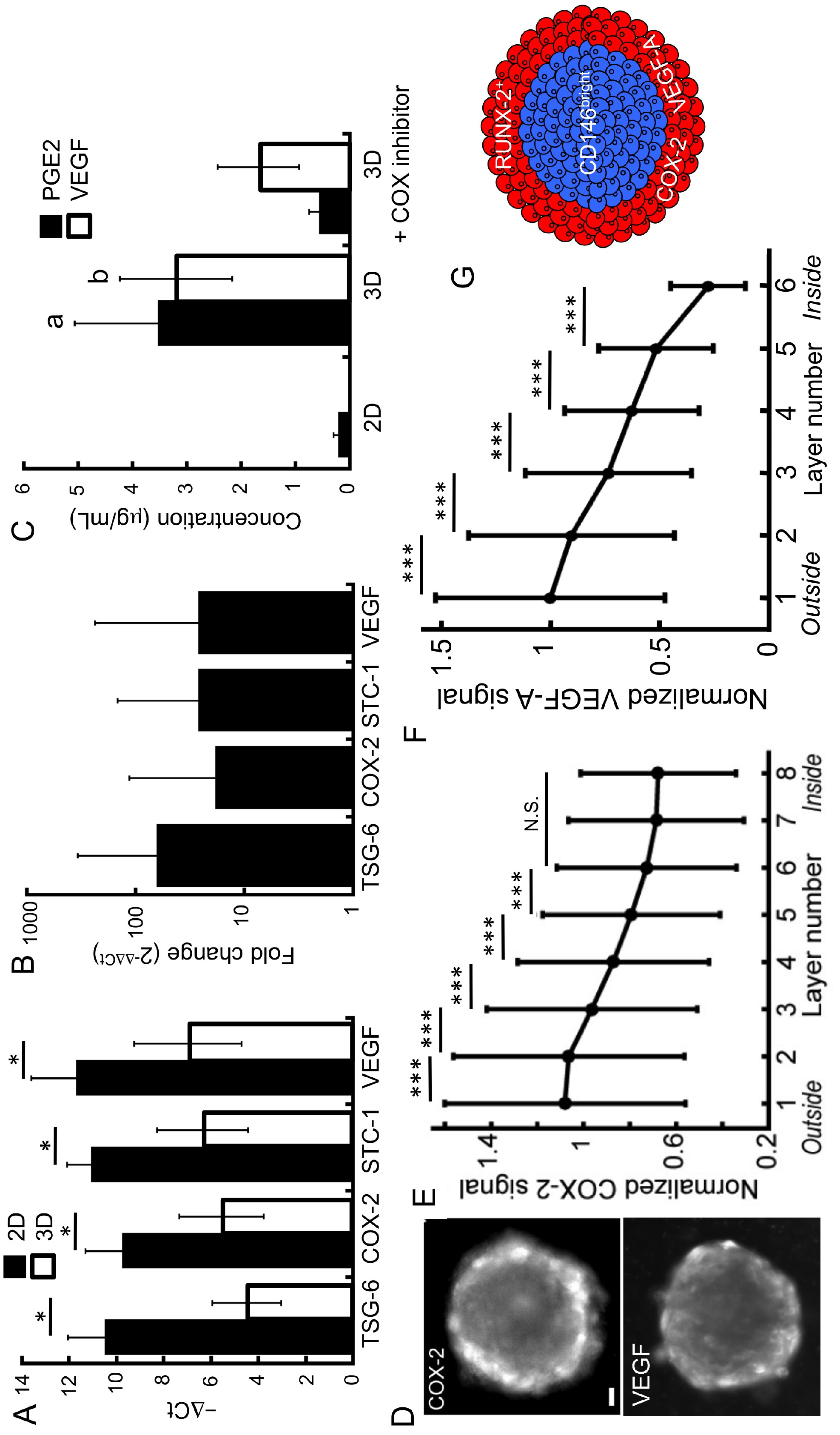
Spatial patterning of the biological functions of hMSCs in 3D. (A-B) RT-qPCR analysis of the relative TSG-6, COX-2, STC-1 and VEGF expression to GADPH (ΔCt, D and relative RNA expression, E), in 3D and in the 2D populations (n_3D_=3, n_2D_=3). (C) Quantification by ELISA of the PGE-2 and VEGF secreted by hMSCs cultivated in 2D, as MBs or as MBs treated with indomethacin (n_chips_ = 3; n_2D_ = 3). (D) (D-E) Representative image (D) and quantitative analysis (E) of COX-2 (N_chips_ = 13, n_MBs_ = 2,936) and VEGF (N_chips_ = 3, n_MBs_ = 413) staining within the layers of the MBs, error bars represent the standard deviation) within the cell layers of the MBs. (E-H) of *: p < 0.05; **: p < 0.01; ***: p < 0.001.

To further interrogate the link between the COX-2 and the VEGF producing cells, their location was analyzed by quantitative image analysis at a layer-by-layer resolution. These measurements showed significantly higher levels of COX-2 in the first two layers, compared with the successive layers of the MBs (Fig. 5.D,E), with a continuous decrease of about 40% of the COX-2 signal between the edge and the core. Similar observations were made with VEGF (Fig. 5.D,F), which demonstrated that cells at the boundaries of the MBs expressed both COX-2 and VEGF (Fig. 5.G). Taken with the measurements of Fig. 5.C, these results imply that COX-2 acts as an upstream regulator of PGE-2 and VEGF secretion. Of note, oxygen deprivation was unlikely to occur within the center of the MBs, since no hypoxic area was detected through the whole MBs (Fig. S5). Consequently, it is unlikely that HIF signaling mediates the increase VEGF expression at the boundaries of the MBs.

Since variations of COX-2 and *adherens* junctions distribution are co-localized within the MBs (Fig. 4.E-G and Fig. 5.D-E), the results point to a link between the quality of cell-cell interactions and the spatial distribution of the COX-2^high^ cells in 3D. The mechanisms leading to the spatial patterning of COX-2 expression in the MBs were therefore explored using inhibitors of the signaling pathways related to anti-inflammatory molecule production and of the molecular pathways regulating the structural organization (Table S1): *(i)* QNZ that inhibits NF-κB, a critical transcription factor regulating the level of COX-2 expression^45^ *(ii)* DAPT that inhibits the canonical Notch pathway, modulating cell-cell interactions and several differentiation pathways. *(iii)* Y-27632 (Y27) that inhibits ROCK involved in the bundling of F-actin (i.e. formation of stress fibers) to assess the role of acto-myosin organization, and *(iv)* Cytochalasin D (CytoD) to inhibit the polymerization of actin monomers.

While the addition of DAPT had virtually no effect on the ability of the cells to form MBs, Y27 led to MBs with more rounded cells, and both QNZ and CytoD strongly interfered with the MB formation process (Fig. 6.A-C). The results indicate that NF-κB activation and the promotion of actin polymerization are critical signaling steps initiating the process of MB formation by hMSCs.

**Figure 6.**
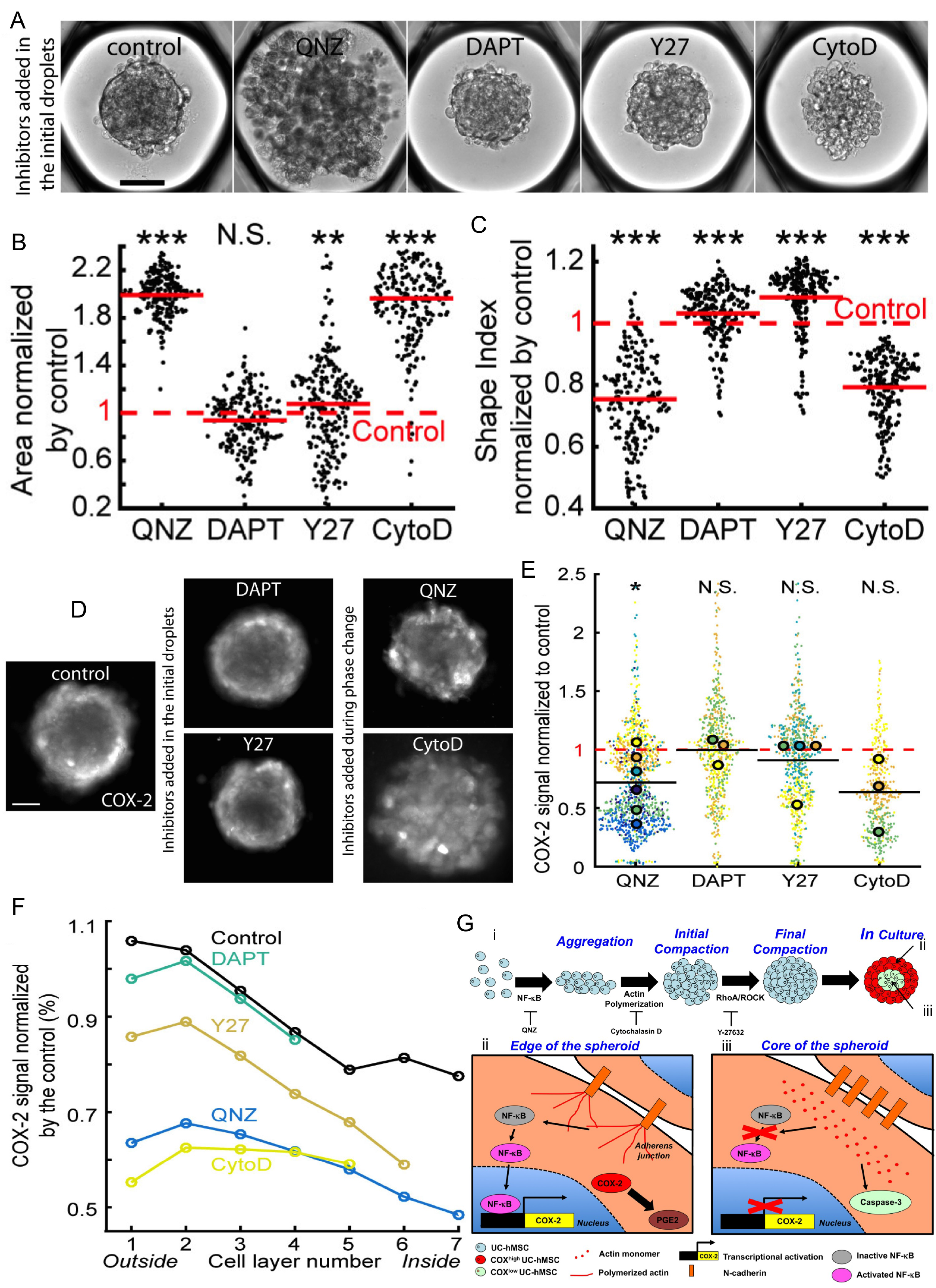
Molecular mechanisms regulating the local level of COX-2 expression in mesenchymal bodies. (A) Representative images of MBs formed 1 day after the droplet loading, scale bar is 100 μm. Inhibitors are added to the culture medium prior to MB formation. (B-C) Quantitative analysis the aggregates projected area (B) and shape index (C) in the presence of the different inhibitors. Red lines represent the mean value for each condition. (D-E) Representative images (D, contrast is adjusted individually for a better visualization of the pattern, scale bar is 100 μm) and quantitative analysis (E) of the COX-2 fluorescent signal intensity normalized by the control value, with the different inhibitors. For these longer culturing times, QNZ and CytoD are only added during the phase change to allow MB formation. Small dots represent one MB. Large dots represent the average normalized COX-2 fluorescent signal per chip. Each color corresponds to a specific chip. *: p < 0.05; N.S. : non-significant. (F) Estimation of inhibitor effect in the cell layers with the COX-2 signal normalized by the control value. Control: N_chips_ = 11, n_MBs_ = 2,204; QNZ: N_chips_ = 6, n_MBs_ = 1,215; DAPT: N_chips_ = 3, n_MBs_ = 658; Y-27: N_chips_ = 4, n_MBs_ = 709; CytoD: N_chips_ = 3, n_MBs_ = 459. *: p < 0.05; **: p < 0.01; ***: p < 0.001; N.S.: non significant. (G) Proposed mechanisms regulating mesenchymal bodies formation and the patterning of their biological functions. (i) Regulation of the formation of mesenchymal bodies. (ii-iii) Spatial patterning of hMSC biological properties within mesenchymal bodies.

To assess the role of NF-κB and actin polymerization in the pattern and the level of COX-2 expression in the MBs, QNZ and CytoD were added one day after the cell seeding, once the MBs were completely formed. In contrast, Y27 and DAPT were included in the initial droplets and maintained in the culture medium for the whole culture period. Typical images showing the COX-2 signal in these different conditions are shown in Fig. 6.D (see also Fig. S6 for quantification of the individual experiments). Of note, none of the inhibitors had an effect on Casp3 activation indicating that they do not induce apoptosis (Fig. S7). The levels of COX-2 expression in MBs, after three days in culture, were significantly reduced with QNZ, also decreasing after the addition of CytoD (Fig. 6.E). By contrast, Y27 and DAPT had no effect on the levels of COX-2 expression. As a consequence, the results demonstrate that a sustained NF-κB activity after MB formation is required to promote COX-2 expression. Moreover, the induction of actin polymerization in MBs constitutes a mandatory step to initiate COX-2 production.

To get a deeper understanding on the local regulation of these signaling pathway, we analyzed at the single cell resolution the distribution of COX-2 within the MBs. The spatial mapping revealed that the COX-2 fluorescence intensity was mostly attenuated at the edge of the MBs treated with CytoD and QNZ, while more limited change in the pattern of its expression was observed in the presence of Y27 and even less so with DAPT (Fig. 6.F). Consequently, the results revealed a strong link between cell phenotype, the capability to form functional *adherens* junctions and the local regulation of NF-κB and actin polymerization leading to the increased expression of PGE-2 and VEGF that are mediated by COX-2 in 3D (Fig. 6.G). Altogether the results indicate that in 3D cell aggregates the spatial organization determines the specific activation of signaling pathways resulting in local functional heterogeneity.

## 3. Discussion

Understanding the mechanisms of formation and the spatial tissue patterning within organoids requires a characterization at single cell level in 3D. In this study, we used a novel microfluidic and epi-fluorescence imaging technology to obtain a precise quantitative mapping of the structure, the position and the link with individual cell functions within MBs. The image analysis provided quantitative data that were resolved on the scale of the individual cells, yielding measurements on 700,000 cells *in situ* within over 10,000 MBs. A Voronoi segmentation was used to categorize the cells into concentric layers, starting from the edge of the MBs and ending with the cells in the central region^27,28^, which allowed us to measure variations in the structural organization and in the protein expressions on a layer-by-layer basis within the 3D cultures.

The MBs were found to organize into a core region of undifferentiated cells, surrounded by a shell of committed cells. This hierarchical organization results from the spatial segregation of an initially heterogeneous population, as is generally the case for populations of pluripotent and somatic stem cells^2,3,4^. The process of aggregation of hMSCs obtained within a few hours takes place through different stages (Fig. 6.G): the first steps of the aggregation of MBs are mediated by N-cadherin interactions. In parallel NF-κB signaling is activated, promoting cell survival by preventing anoikis of suspended cells^46,47^. At later stages, the formation of polymerized F-actin and, to a lesser extent, stress fibers mediate the MB compaction, mainly at the edge of the MBs where the cellular commitment helps the stabilization of *adherens* junctions. The formation of *adherens* junctions facilitates the cohesion of the 3D structure, probably through to the enhanced β-catenin availability in the CD146^-^/RUNX-2+ cells, which is recruited in the CCC complexes of the *adherens* junctions.

A functional consequence of this hierarchical segregation is an increase of endocrine activity of the cells located at the boundaries of the MBs. Indeed, COX-2 expression is increased in the outer layers of the MBs, which also contain more functional *adherens* junctions as well as a sustained NF-κB activity in this region. The promoter of *COX-2* contains RUNX-2 and NF-κB cis-acting elements^48,49^. While RUNX-2 is required for COX-2 expression in mesenchymal cells, its level of expression does not regulate the levels of COX-2^48,49^. The increased COX-2 expression is in turn due to the unbundled form of F-actin (i.e. a more relaxed form of actin, in comparison to the dense stress fibers observed in 2D) near the edge of the MBs, which was reported to sustain NF-κB activity^50^ and to downregulate COX-2 transcriptional repressors^51,52^. Therefore NF-κB has a high activity in the outer layers of the MB, where it locally promotes COX-2 expression.

These results show that the 3D culture format is fundamental to understand the mesenchymal cell behavior *in vivo*, since we found that the expression of key bone regulatory molecules is spatially regulated as a function of the structural organization of the MBs. The 3D structure obtained here mimics the conditions found at the initial steps of intramenbranous ossification that occurs after mesenchymal condensation *(i.e*. no chondrogenic intermediate was found in the MBs). In the developing calvaria, the most undifferentiated mesenchymal cells (e.g. Sca-1^+^/RUNX2^-^ cells) are located in the intrasutural mesenchyme, which is surrounded by an osteogenic front containing more committed cells (e.g. Sca-1^-^/RUNX-2^+^ cells)^18,53^. Similarly, we observed that undifferentiated hMSCs (i.e. CD146^+^/RUNX-2^-^ hMSCs) were surrounded by osteogenically committed cells (i.e. CD146^-^/RUNX-2^+^ hMSCs), which also co-expressed pro-osteogenic molecules, namely COX-2 and its downstream targets, PGE2 and VEGF. The present study suggests that such *in vitro* models can be used to understand the early steps of bone formation and the emergence of bone displasias.

This study also reveals the importance of resolving spatial heterogeneities within organoids to link cell structural organization and their functional properties. This data-driven approach of combining high throughput 3D culture and multiscale cytometry^21^ on complex biological models can be applied for getting a better understanding of the equilibria that determine the structure and the function of cells within multicellular tumor spheroids, embryoid bodies, or organoids.

## Acknowledgements

Caroline Frot is gratefully acknowledged for her help with the microfabrication. The research leading to these results received funding from the European Research Council (ERC) Grant Agreement 278248 Multicell.

## Author Contributions

SS, CNB, and AC conceived the experiments. SS performed the experiments. RFXT wrote the image processing code and performed image analysis. RFXT, SS, GA and AB performed image and data analyses. SS, CNB, AC discussed the results and wrote the manuscript. All authors discussed the manuscript.

## METHODS

### Human Umbilical Cord Derived Mesenchymal Stromal Cell Culture

Human mesenchymal stromal cells derived from the Wharton’s jelly of umbilical cord (hMSCs) (ATCC® PCS-500-010, American Type Culture Collection, LGC, Molsheim, France) were obtained at passage 2. Three different lots of hMSCs were used in this study (lot # 60971574, lot # 63739206 and lot # 63516504), which showed consistent results for COX-2 and CD146 distribution. hMSCs from the different lots were certified for being CD29-, CD44-, CD73-, CD90-, CD105-, CD166- positive (more than 98 % of the population is positive for these markers) and CD14-, CD31-, CD34-, CD45- negative (less than 0.6 % of the population is positive for these markers) and to differentiate into adipocytes, chondrocytes, and osteocytes (ATCC, certificate of analysis). hMSCs were maintained in T-175 cm^2^ flasks (Corning, France) and cultivated in a standard CO_2_ incubator (Binder, Tuttlingen, Germany). The culture medium was composed of Alpha Modified Eagle’s medium (α-MEM) (Gibco, Life Technologies, Saint Aubin, France) supplemented with 10 % (v/v) fetal bovine serum (Gibco) and 1 % (v/v) penicilin-streptamicine (Gibco). The cells were seeded at 5.10^3^ cells/cm^2^, sub-cultivated every week, and the medium was refreshed every 2 days. hMSCs at passage 2 were first expanded until passage 4 (for about 5-6 populations doublings, PDs), then cryopreserved in 90 % (v/v) FBS / 10 (v/v) % DMSO and stored in a liquid nitrogen tank. The experiments were carried out with hMSCs at passage 4 to 11 (about 24-35 PDs, after passage 2).

### Surface Markers Staining, Analysis and Sorting by Flow Cytometry

hMSCs were harvested by scrapping or trypsinization from T-175 cm^2^ flasks. Then, the cells were incubated in staining buffer (2% FBS in PBS), stained with a mouse anti-human CD146-Alexa Fluor^®^647 (Clone P1-H12, BD Bioscience), a mouse anti-human CD31-Alexa Fluor^®^488 (BD Bioscience, San Jose, CA) antibody, a mouse anti-human CD105-Alexa Fluor^®^ 647 (BD Bioscience, San Jose, CA), a mouse anti-human CD 90-FITC and a mouse anti-human CD73-APC (Miltenyi Biotec, Germany).

The percentages of CD73, CD90, CD105, CD146 and CD31 positive cells were analyzed using a FACS LSRFortessa (BD Bioscience, San Jose, CA). To validate the specificity of the antibody staining, the distributions of fluorescently labeled cells were compared to cells stained with isotype controls: mouse IgG1, k-PE-Cy™5 (clone MOPC-21, BD Bioscience) and mouse IgG2a K isotype control FITC (BD Bioscience, San Jose, CA). Alternatively, hMSCs were sorted based on their level of expression of CD146 or their size (FSC and SSC) using a FACSAria III (BD Bioscience, San Jose, CA).

### Adipogenic and Osteogenic Differentiation

To induce adipogenic differentiation, UC-hMSCs were seeded at 1.10^4^ cells/cm^2^ in culture medium. The day after, the culture medium was switched to StemPro® Adipogenesis Differentiation medium (Life Technologies) supplemented with 10 μM Rosiglitazone (Sigma-Aldrich) for two weeks. To visualize the differentiated adipocytes, the cells were stained with Oil-red O (Sigma-Aldrich). As a control, UC-hMSCs were maintained in culture medium for two weeks and stained with Oil-red O as above.

To induce osteogenic differentiation, UC-hMSCs were seeded at 5.10^3^ cells/cm^2^ in culture medium. The day after, the culture medium was switched to StemPro® Osteogenesis Differentiation medium (Life Technologies) supplemented with 2 nM BMP-2 (Sigma-Aldrich) for two weeks. To visualize the differentiated osteoblasts, the cells were stained with Alizarin Red S (Sigma-Aldrich). As a control, UC-hMSCs were maintained in culture medium for two weeks and stained with Alizarin Red S, as above.

### Microfabrication

Standard dry film soft lithography was used for the flow-focusing device (top of the chip) fabrication, while a specific method for the fabrication of the anchors (bottom of the chip) was developed. For the first part, up to five layers of dry film photoresist consisting of 50 μm Eternal Laminar E8020, 33 μm Eternal Laminar E8013 (Eternal Materials, Taiwan) and 15 μm Alpho NIT215 (Nichigo-Morton, Japan) negative films were successively laminated using an office laminator (PEAK pro PS320) at a temperature of 100 °C until the desired channel height, either 135, 150, 165 or 200 μm, was reached. The photoresist film was then exposed to UV (LightningCure, Hamamatsu, Japan) through a photomask of the junction, channels and the culture chamber boundaries. The masters were revealed after washing in a 1 % (w/w) K_2_CO_3_ solution (Sigma-Aldrich). For the anchors fabrication, the molds were designed with RhinoCAM software (MecSoft Corporation, LA, USA) and were fabricated by micro-milling a brass plate (CNCMini-Mill/GX, Minitech Machinery, Norcross, USA). The topography of the molds and masters were measured using an optical profilometer (VeecoWyco NT1100, Veeco, Mannheim, Germany).

For the fabrication of the top of the chip, poly(dimethylsiloxane) (PDMS, SYLGARD 184, Dow Corning, 1:10 (w/w) ratio of curing agent to bulk material) was poured over the master and cured for 2 h at 70 °C. For the fabrication of bottom of the chip, the molds for the anchors were covered with PDMS. Then, a glass slide was immersed into uncured PDMS, above the anchors. The mold was finally heated on a hot plate at 180 °C for 15 min. The top and the bottom of chip were sealed after plasma treatment (Harrick, Ithaca, USA). The chips were filled 3 times with Novec Surface Modifer (3M, Paris, France), a fluoropolymer coating agent, for 30 min at 110 °C on a hot plate.

### Formation of Mesenchymal Bodies on Chip

hMSCs were harvested with trypLE™ at 60 – 70 % confluence and a solution containing 6.10^5^ cells in 70 μL medium were mixed with a 30 μL 3 % (w/v) liquid low melting agarose solution (i.e. stored at 37 °C) (Sigma-Aldrich, Saint Quentin Fallavier, France) diluted in culture medium containing 50 μg/mL Gentamicin (Sigma-Aldrich) (1:3 v/v), resulting in a 100 μL solution of 6.10^6^ cells/mL in 0.9 % (w/v) agarose.

hMSCs and agarose were loaded into a 100 μL glass syringe (SGE, Analytical Science, France), while Fluorinert® FC-40 oil (3M, Paris, France) containing 1 % (w/w) PEG-di-Krytox surfactant (RAN Biotechnologies, Bervely, USA) was loaded into a 1 mL and 2.5 mL glass syringes (SGE, Analytical Science). Droplets of cell-liquid agarose were generated in the FC-40 containing PEG-di-Krytox, at the flow-focusing junction, by controlling the flow rates using syringe pumps (neMESYS Low Pressure Syringe Pump, Cetoni GmbH, Korbussen, Germany) (Table S1). After complete loading, the chips were immersed in PBS and the cells were allowed to settle down and to organize as mesenchymal bodies (MBs) overnight in the CO_2_ incubator. Then, the agarose was gelled at 4 °C for 30 min, after which the PEG-di-Krytox was extensively washed in flushing pure FC-40 in the culture chamber. After washing, cell culture medium was injected to replace the FC-40. All flow rates are indicated in Table S1. Further operations were allowed by gelling the agarose in the droplets, such that the resulting beads were retained mechanically in the traps rather than by capillary forces (Fig. 2.G). This step allowed the exchange of the oil surrounding the droplets by an aqueous solution, for example in order to bring fresh medium for long term culture, chemical stimuli, or the different solutions required for cell staining.

### Live Analysis of MB Formation

For the live imaging of the MB formation, the chips were immersed in PBS, and then were incubated for 24 hours in a microscope incubator equipped with temperature, CO_2_ and hygrometry controllers (Okolab, Pozzuoli, Italy). The cells were imaged every 20 min.

### Immunocytochemistry

2D cultures or MBs were washed in PBS, and incubated with a 5 μM Nucview™ 488 caspase-3 substrate (Interchim, Montluçon, France) diluted in PBS. After washing with PBS, hMSCs were fixed with a 4 % (w/v) PFA (Alpha Aesar, Heysham, UK) for 30 min and permeabilized with 0.2-0.5 % (v/v) Triton X-100 (Sigma-Aldrich) for 5 min. The samples were blocked with 5 % (v/v) FBS in PBS for 30 min and incubated with a rabbit polyclonal anti-COX-2 primary antibody (ab15191, Abcam, Cambridge, UK) diluted at 1:100 in 1 % (v/v) FBS for 4 h. After washing with PBS, the samples were incubated with an Alexa Fluor^®^594 conjugate goat polyclonal anti-rabbit IgG secondary antibody (A-11012, Life Technologies, Saint Aubin, France) diluted at 1:100 in 1 % (v/v) FBS, for 90 min. Finally, the cells were counterstained with 0.2 μM DAPI for 5 min (Sigma-Aldrich), and then washed with PBS.

The same protocol was used for the staining of VEGF-A expressing cells using a rabbit anti-human VEGF-A monoclonal antibody (ab52917, Abcam, Cambridge, UK), which was revealed using the same secondary antibody as above. RUNX-2 positive cells were similarly stained using a mouse anti-human RUNX-2 monoclonal antibody (ab76956, Abcam, Cambridge, UK), which was revealed using an Alexa Fluor^®^488 goat anti-mouse IgG2a secondary antibody (A-21131, Life Technologies, Saint Aubin, France), both diluted at 1:100 in 1 % (v/v) FBS.

### Detection of Hypoxia Within the MBs

To measure potential induction of hypoxia within the core of the MBs, the cells were stained with Image-iT™ Red Hypoxia Reagent (Invitrogen) for 3 hours, then imaged using a fluorescent microscope. As a positive control, the chips containing the MBs were immersed into PBS and incubated overnight in an incubator set at 37°C under 3% O_2_/5% CO_2_, and finally imaged as above.

### Inhibition of Molecular Pathways Regulating Properties of hMSCs in MBs

To interrogate the contribution of signaling related to anti-inflammatory molecules production (COX-2, NF-κB), or molecular pathways regulated by the cell structural organization (Notch, ROCK and F-actin), several small molecules inducing their inhibition were added to the culture medium (Table S1). For all the conditions, the final concentration of DMSO was below 0.1 % (v/v) in the culture medium.

### Viability Assay

The cell viability was assessed using LIVE/DEAD^®^ staining kit (Molecular Probes, Life Technologies). The MBs were incubated for 30 min in PBS containing 1 μM calcein AM and 2 μM ethidium homodimer-1 (EthD1), in flushing 100 μL of the solution. The samples were then washed with PBS and imaged under a motorized fluorescent microscope (Nikon, France).

### N-Cadherin Immunostaining

For the detection of the functional forms of N-cadherins (i.e. the N-cadherins closely linked to the actin network, which are PFA insoluble), the MBs were fixed with a 4 % (w/v) PFA (Alpha Aesar, Heysham, UK) for 30 min and permeabilized with 0.2-0.5 % (v/v) Triton X-100 (Sigma-Aldrich) for 5 min. Alternatively, the aggregates were incubated for 5 min with 100 % cold methanol followed by 1 min with cold acetone, for the detection of total N-cadherins (i.e. the PFA soluble and insoluble forms).

Then, the samples were blocked with 5 % (v/v) FBS in PBS for 30 min and incubated with a Rabbit polyclonal anti N-cadherin primary antibody (ab18203, Abcam, Cambridge, UK) diluted at 1:100 in 1 % (v/v) FBS for 4 h. After washing with PBS, the samples were incubated with an Alexa Fluor^®^594 conjugate goat polyclonal anti-rabbit IgG secondary antibody (A-11012, Life Technologies, Saint Aubin, France) diluted at 1:100 in 1 % (v/v) FBS, for 90 min. Finally, the cells were counterstained with 0.2 μM DAPI for 5 min (Sigma-Aldrich), and then washed with PBS.

### F-Actin Staining

For the quantification of the polymerized form of actin (F-actin), the MBs were first fixed with a 4 % (w/v) PFA (Alpha Aesar, Heysham, UK) for 30 min and permeabilized with 0.2-0.5 % (v/v) Triton X-100 (Sigma-Aldrich) for 5 min. The samples were then blocked with a 5 % (v/v) FBS solution and incubated for 90 min in a 1/100 phalloidin-Alexa^®^594 (Life Technologies) diluted in a 1 % (v/v) FBS solution. The cells were then counterstained with 0. 2 μM DAPI for 5 min (Sigma-Aldrich), and then washed with PBS.

### Validation of the of the Fluorescent Signal Patterns

To ensure the specificity of the antibody to COX-2 and N-cadherin, control UC-hMSCs were permeablized, fixed and incubated only with the secondary antibody (Alexa Fluor® 594 conjugate goat polyclonal anti-rabbit IgG), as above. The absence of fluorescent signal indicated the specific staining for intracellular COX-2 and N-cadherin.

Next, in order to validate that the distribution of the fluorescent intensity was not related to any antibody diffusion limitation, the MBs were fixed and permabilized as above. For this assay, the MBs were not subjected to any blocking buffer. The cells were incubated for 90 min with the Alexa Fluor® 594 conjugate goat polyclonal anti-rabbit IgG secondary antibody (A-11012, Life Technologies, Saint Aubin, France) diluted at 1:100 in 1 % (v/v) FBS. Then, the cells were counterstained for DAPI as above. Finally the MBs were collected from the chip, deposed on a glass slide and imaged.

For the analysis of COX-2 expression by flow cytometry, the total MBs were recovered from the chip. The MBs were then trypsinized and triturated to obtain single cell suspension. UC-hMSCs were stained for COX-2 as above. The percentage of COX-2 positive cells was quantified on 5.10^3^ dissociated UC-hMSCs using a Guava^®^ easyCyte Flow Cytometer (Merck Millipore, Guyancourt, France). The results were compared to the fluorescent intensity distribution obtained by image analysis.

### Clearing MBs derived from hMSCs

To interrogate the influence of the MBs opacity in the COX-2 and N-cadherin fluorescent signals, the samples were treated by the Clear(T2) method, after immunostaining^54^. Briefly, the MBs were incubated for 10 min in 25 % (v/v) formamide / 10 % (w/v) PEG (Sigma-Aldrich), then for 5 min in 50 % (v/v) formamide / 20 % (w/v) PEG, and finally for 60 min in 50 % (v/v) formamide / 20 % (w/v) PEG, prior to their imaging. The fluorescent signal distribution was compared to non-cleared samples.

### Cryosections

The MBs were collected from the chip, and then fixed using PFA as above. The MBs were incubated overnight in a 30% sucrose solution at 4ºC. Then, the sucrose solution was exchanged to O.C.T. medium (Optimal Cutting Temperature, Tissue Tek) in inclusion molds, which were slowly cooled down using dry ice in ethanol. The molds were then placed at -80 ºC. The day of the experiments, the O.C.T. blocks were cut at 7 μm using a cryostat (CM 3050S, Leica). The cryosections were placed on glass slides (SuperFrost Plus Adhesion, Fisher Scientific), dried at 37°C and rehydrated using PBS. The cryosections were permeabilized and stained for COX-2 as above. The slides were finally mounted in mounting medium containing DAPI (Fluoromount-G, Invitrogen^TM^).

### Competitive Enzyme-Linked Immunosorbent Assay for PGE-2

The culture supernatants of 6 well plates were collected, while the total medium content of the chip was recovered by flushing the culture chamber with pure oil. A PGE-2 human ELISA kit (ab133055, Abcam, Cambridge, UK) was used for the quantification of PGE-2 concentration in the culture supernatant, following the manufacturer instructions. Briefly, a polynomial standard curve of PGE-2 concentration derived from the serial dilution of a PGE-2 standard solution was generated (r^2^ > 0.9). The absorbance was measured using a plate reader (Chameleon, Hidex, Finland).

### Enzyme-Linked Immunosorbent Assay for VEGF-A

A VEGF-A human ELISA kit (Ab119566, Abcam, Cambridge, UK) was used for the quantification of VEGF-A concentration in the culture supernatant of 2D cultures or from the chips. A linear standard curve of VEGF-A concentration derived from the serial dilution of a VEGF-A standard solution was generated (r^2^ > 0.9). The absorbance was measured using a plate reader (Chameleon, Hidex, Finland).

### RT-qPCR Analysis

The total MBs of a 3-day culture period were harvested from the chips, as described above. Alternatively, cells cultured on regular 6 well plates were recovered using trypsin after the same cultivation time; CD146^dim^ and CD146^bright^ isolated cells were immediately treated for RNA extraction after sorting. The total RNA of 1.10^4^ cells were extracted and converted to cDNA using Superscript^TM^ III CellsDirect cDNA synthesis System (18080200, Invitrogen, Life Technologies), following the manufacturer instructions. After cell lysis, a comparable quality of the extracted RNA was observed using a bleach agarose gel and similar RNA purity was obtained by measurement of the optical density at 260 nm and 280 nm using a NanoDrop spectrophotometer (Thermo scientific, Wilmington, DE, USA), between total RNA preparations from 2D and on-chip cultures.

The cDNA were amplified using a GoTaq^®^qPCR master mix (Promega, Charbonnieres, France) or a FastSart Universal SYBR Green master mix (containing Rox) (Roche), and primers (Life technologies, Saint Aubin, France or Eurofins Scientific, France) at the specified melting temperature (Tm) (Table S2), using a Mini-Opticon^TM^ (Bio-Rad) or a Quantstudio 3 (Thermo Scientific) thermocycler. As negative control, water and total RNA served as template for PCR. To validate the specificity of the PCR reaction, the amplicons were analyzed by dissociation curve and subsequent loading on a 2.5 % (w/v) agarose gel and migration at 100 V for 40 min. The PCR products were revealed by ethidium bromide (Sigma-Aldrich) staining, and the gels were imaged using a trans-illuminator. The analysis of the samples non-subjected to reverse transcription (RT^-^) indicated negligible genomic DNA contamination (i.e. < 0.1 %), while no amplification signal was observed for the water template (NTC). The amount of TSG-6, COX-2, STC-1, VEGF-A, RUNX-2, CEBP-α and SOX-9 transcripts was normalized to the endogenous reference (GADPH), and the relative expression to a calibrator (2D cultures) was given by 2^-ΔΔ^Ct^^ calculation. A least five biological replicates of 2D and on-chip cultures were analyzed by at least duplicate measurements. The standard curves for GADPH, TSG-6, COX-2 and STC-1 were performed using a 5 serial dilution of the cDNA templates, and indicated almost 100 % PCR efficiency.

### Image Acquisition and Analysis

The image analysis allowed to perform a multiscale analysis^21^ of the MBs. For each chip, single images of the anchors were acquired automatically with the motorized stage of the microscope. The analysis was conducted on a montage of the detected anchors using a custom Matlab code (R2016a, Mathworks, Natick, MA, USA). Two distinct routines were used: one with a bright field detection, and one for the fluorescence experiments.

For bright field detection was described previously^21^, the cells were detected in each anchor as pixels with high values of the intensity gradient. This allowed for each cell aggregate to compute morphological parameters such as the projected area *A* and the shape index *SI* that quantifies the circularity of an object:

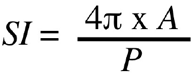

where *P* is the perimeter. Shape index values range from 0 to 1, 1 being assigned for perfect disk.

The MB detection with fluorescent staining (DAPI/Casp3/COX-2, DAPI/Phalloidin, DAPI/N-Cadherin, or LIVE/DEAD^®^) was performed as described previously^21^. First, morphological data was extracted at the MB level, such as the equivalent diameter of the MBs or the *SI*. Also, the mean fluorescent signal of each MB was defined as the subtraction of the local background from the mean raw intensity.

At the cellular level, two different methods were used, both relying on the detection of the nuclei centers with the DAPI fluorescent signal. On the first hand, each cell location could be assigned to a normalized distance from the MB center (r/R) in order to correlate a nuclear fluorescent signal with a position in the MB, as previously described^21^. On the second hand, the cell shapes inside the MBs were approximated by constructing Voronoi diagrams on the detected nuclei centers. Basically, the edges of the Voronoi cells are formed by the perpendicular bisectors of the segments between the neighboring cell centers. These Voronoi cells were used to quantify the cellular cytoplasmic signal (COX-2, F-Actin and N-Cadherin, VEGF and RUNX-2). In details, in order to account for the variability of the cytoplasmic signal across the entire cell (nucleus included), the fluorescent signal of a single cell was defined as the mean signal of 10 % highest pixels of the corresponding Voronoi cell.

Image processing was also used to get quantitative data on 2D cultures, as previously described^21^. Finally, different normalization procedures were chosen in this paper. When an effect was quantified compared to a control condition, the test values were divided by the mean control value and the significance was tested against 1. For some other data, the values were simply normalized by the corresponding mean at the chip level in order to discard the inter-chip variation from the analysis.

### Statistical Analysis

*: p < 0.05; **: p < 0.01; ***: p < 0.001; N.S.: non-significant. Details of each statistical test and p-values can be found in Table S4.

## SUPPLEMENTARY TABLES

**Table 1.**
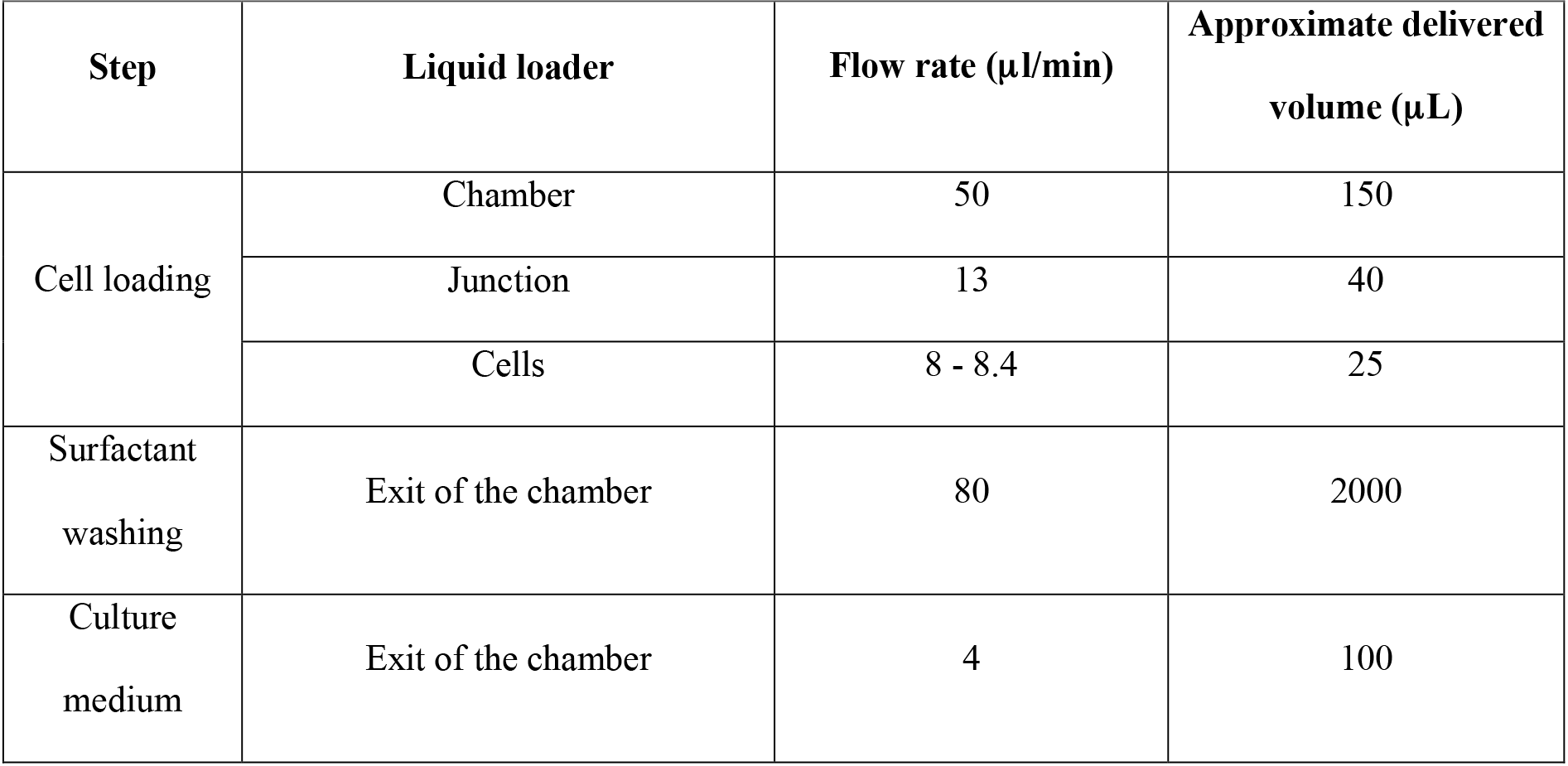
Flow Rate for Cell Loading and Phase Exchange.

| Step | Liquid loader | Flow rate ($\mu\text{l}/\text{min}$) | Approximate delivered volume ($\mu\text{L}$) |
| --- | --- | --- | --- |
| Cell loading | Chamber | 50 | 150 |
|  | Junction | 13 | 40 |
|  | Cells | 8 - 8.4 | 25 |
| Surfactant washing | Exit of the chamber | 80 | 2000 |
| Culture medium | Exit of the chamber | 4 | 100 |

**Table 2.**
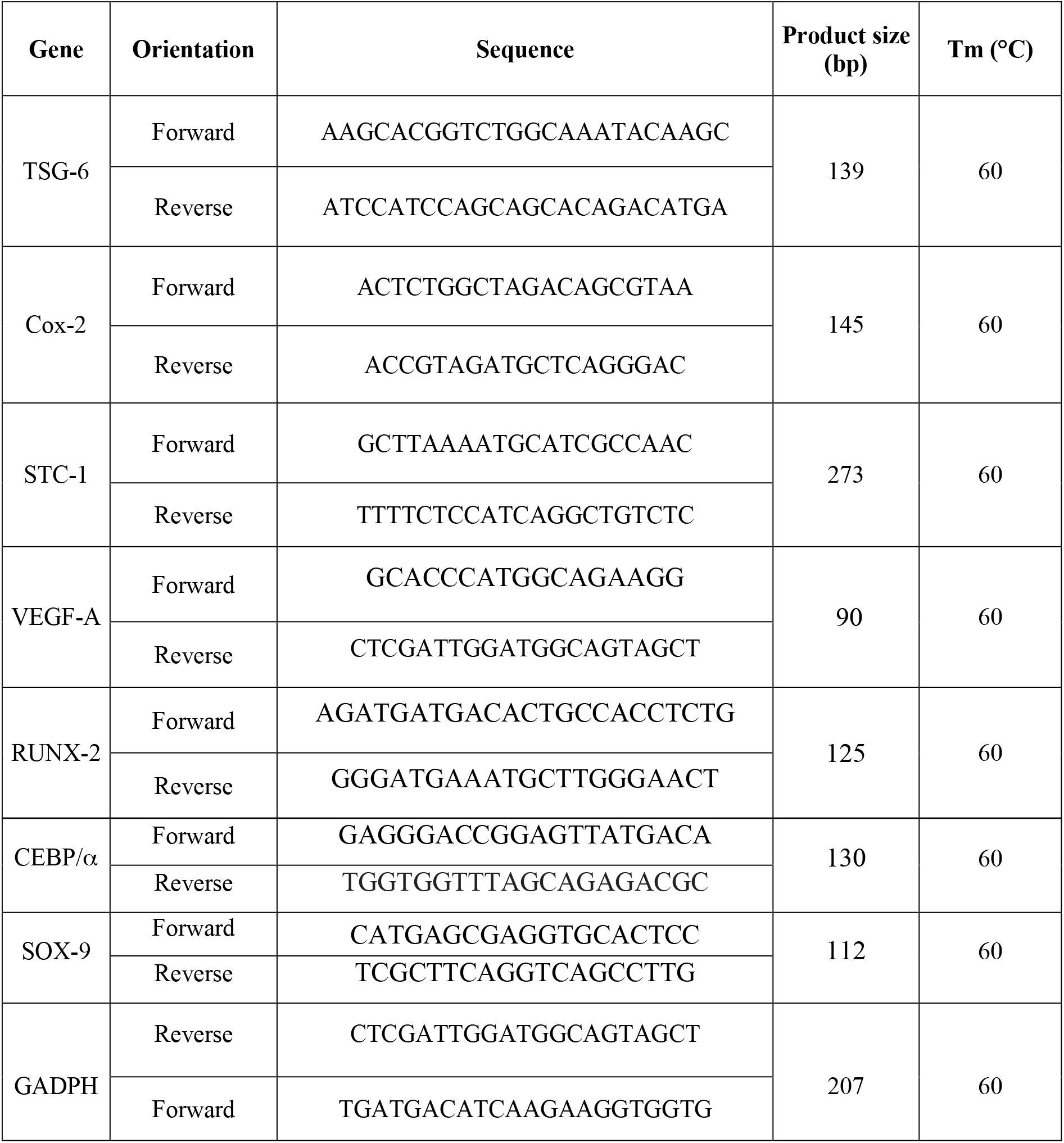
Primer Sequences.

**Table 3.**
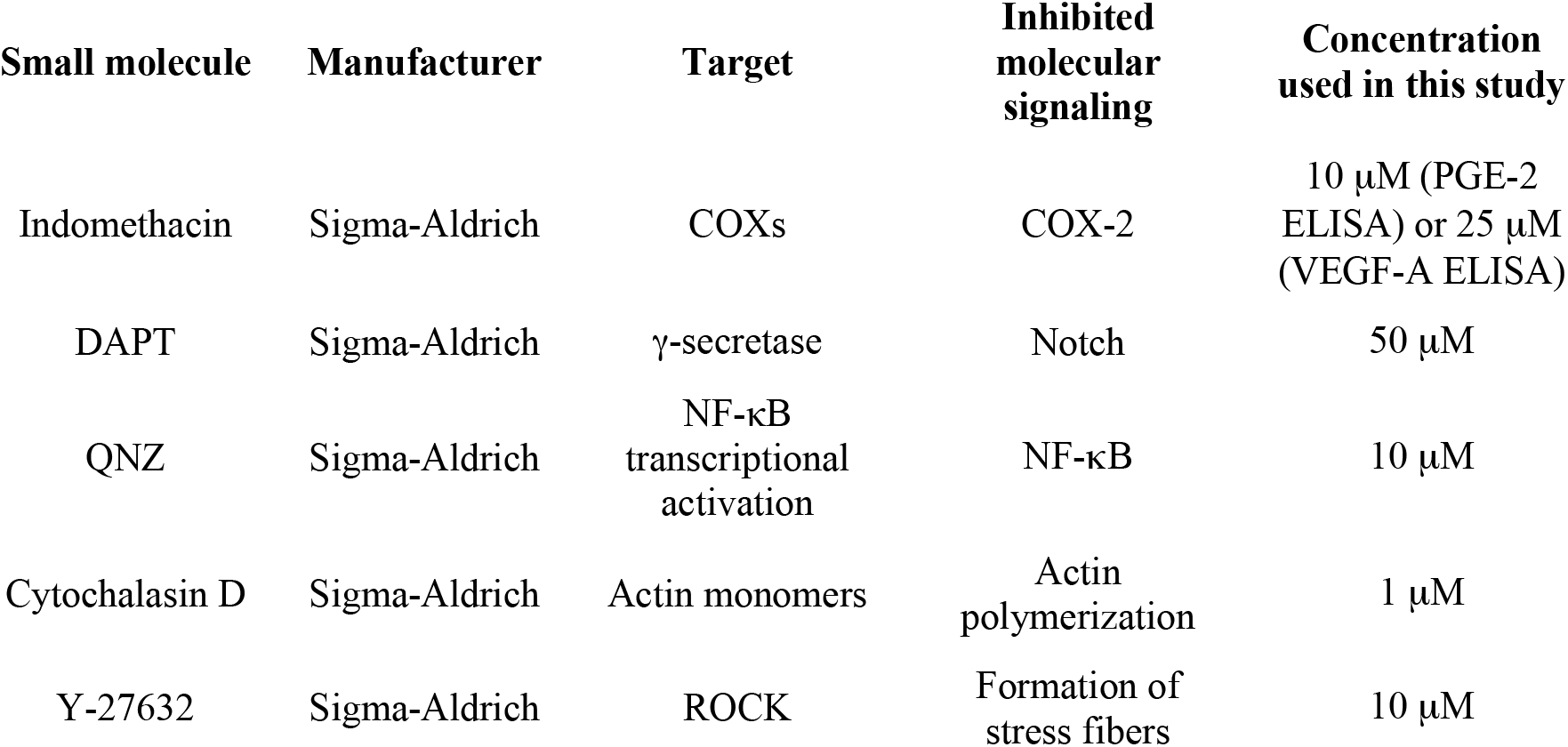
Small Molecules for the Inhibition of Molecular Pathways Regulating hMSC Behavior.

**Table 4.**
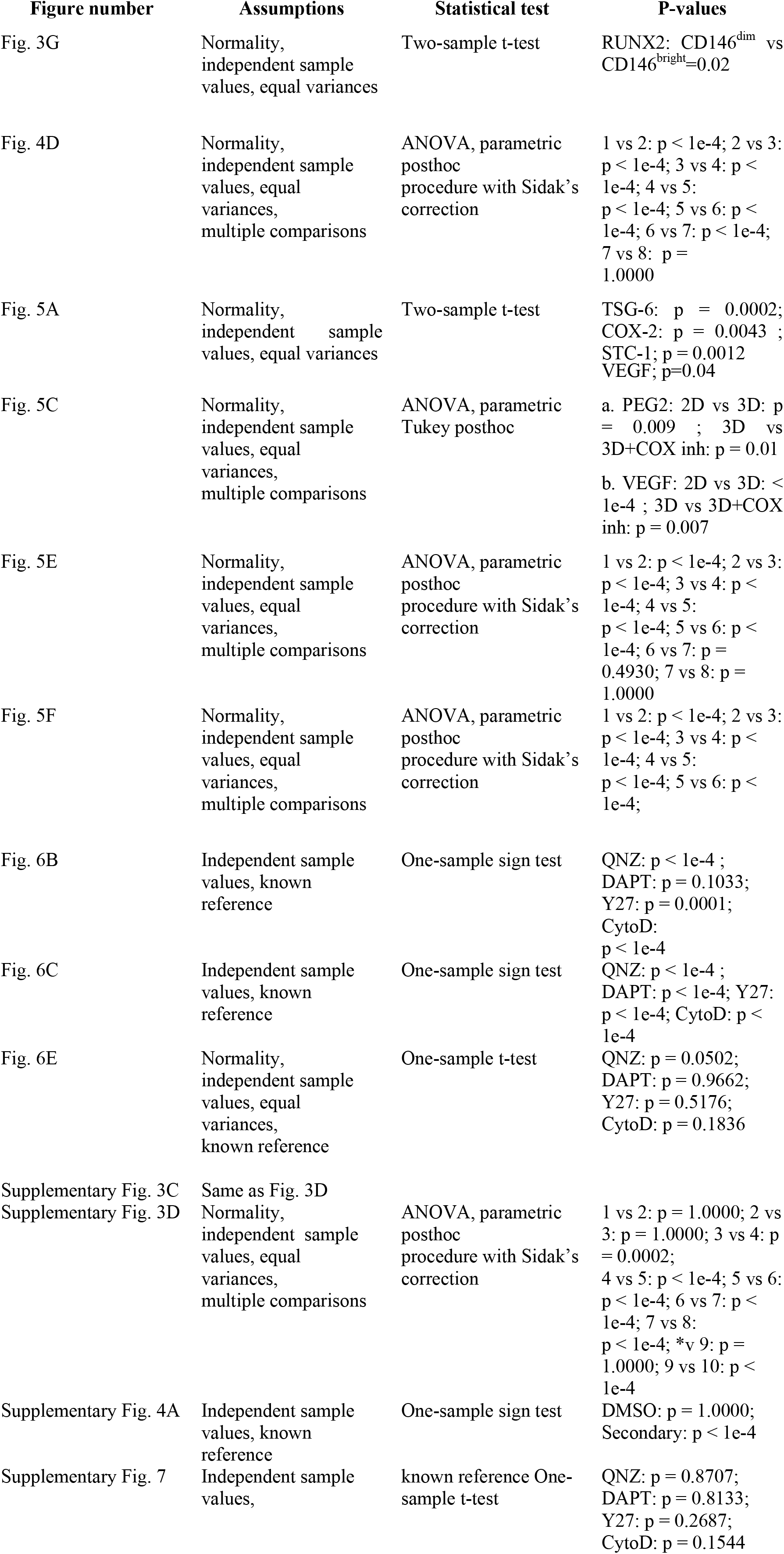
Statistical tests and p-values.

## SUPPLEMENTARY FIGURES

**Supplementary Figure 1.**
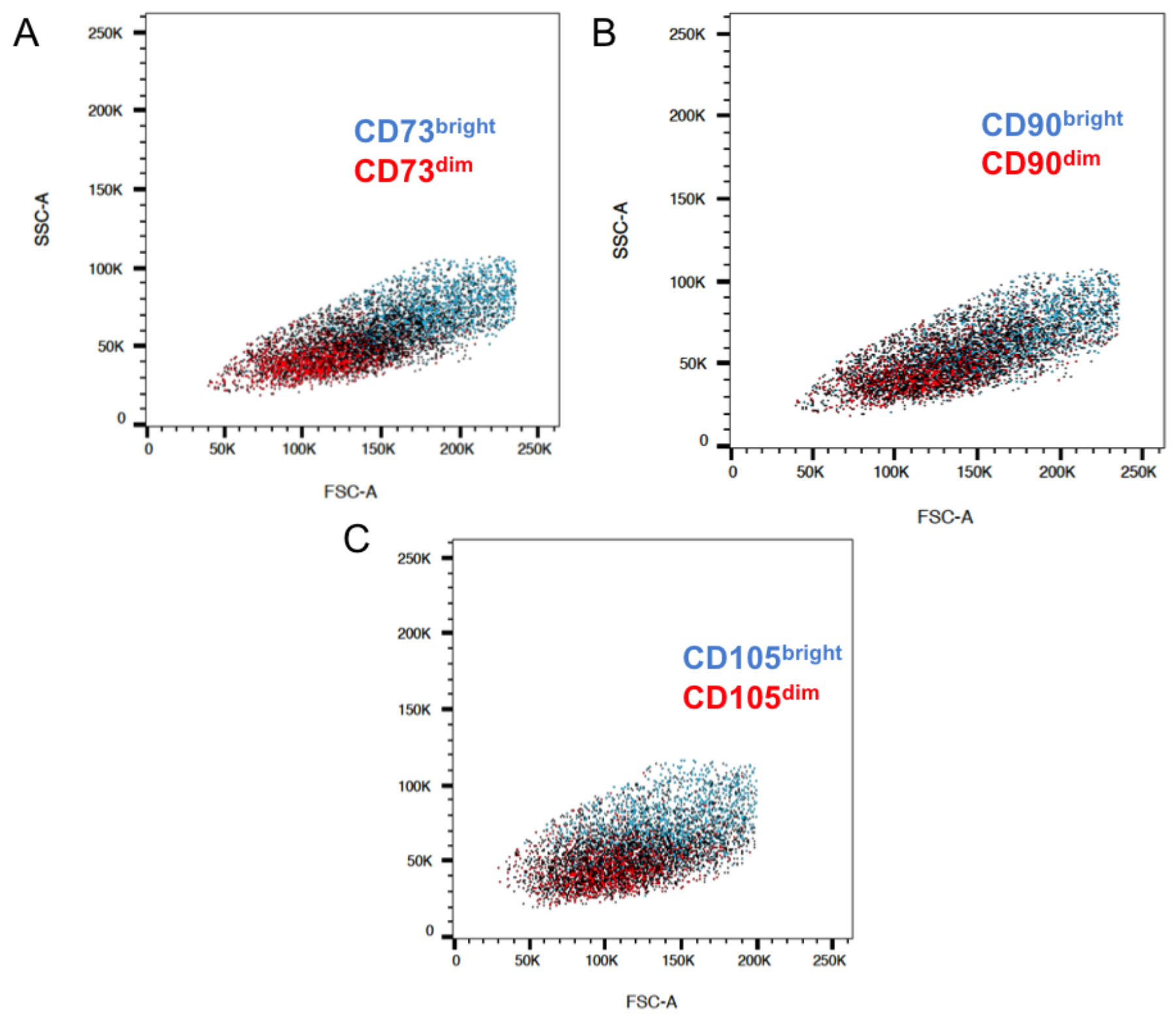
Correlation between cell size (FSC and SSC) and level of CD73 (A), CD90 (B) and CD105 (C) expression.

**Supplementary Figure 2.**
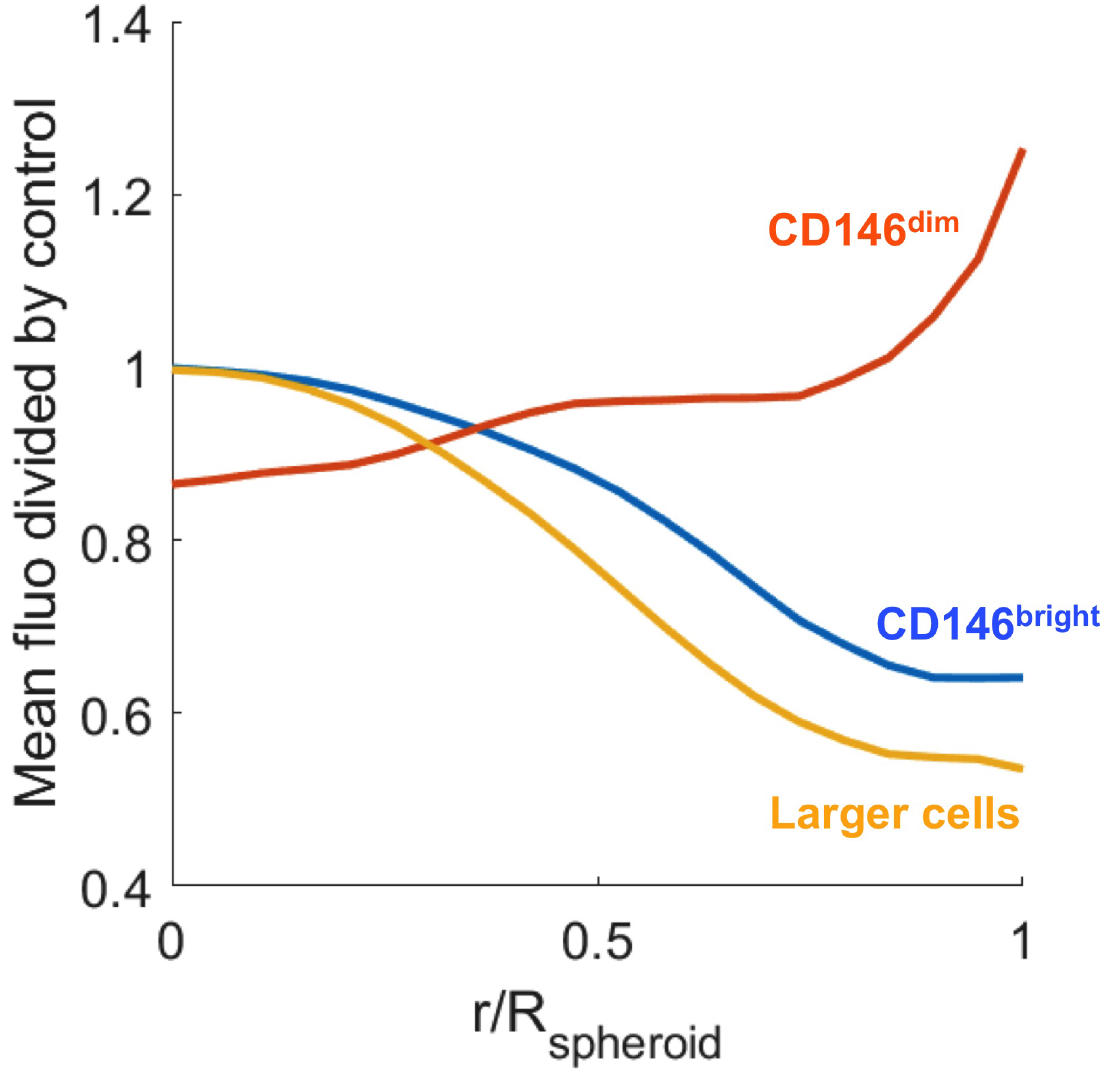
Radial coordinates of CD146^dim^, CD146^bright^ and the larger cells of the hMSC population in MBs.

**Supplementary Figure 3.**
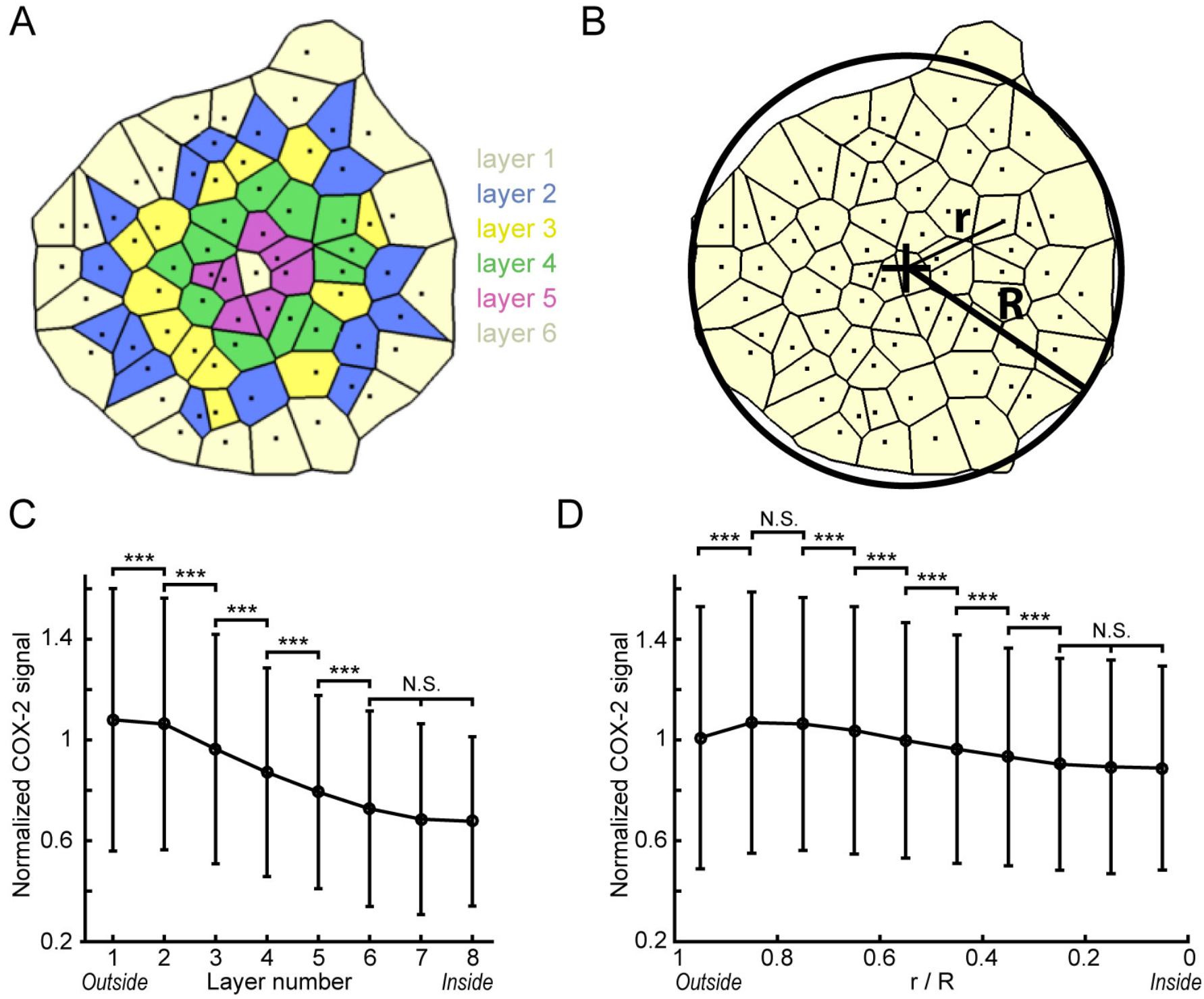
Intra-MB analysis of the COX-2 signal with concentric cell layers. (A-B) Schemes of the same MB where the cell centers are represented by black dots. Black lines show the Voronoi cells. The position of each cell in the MB is described by assigning a cell layer number (A, each color represent a cell layer, layer number 1 being the outermost cell layer) or by computing its normalized distance to the MB center r / R (B, r is the distance between the cell center and the MB center and R is the equivalent radius of the MB). (C-D) Evolution of the normalized cellular COX-2 signal with the cell layer number (C) and the normalized distance to the MB center (D). Black circles and errors bars represent respectively the mean normalized COX-2 signal and the standard deviation of the data. N_chips_ = 13; n_MBs_ = 2,936; n_cells_ = 159,596. The layer analysis gives a much clearer trend with a higher COX-2 signal in the layers close to the MB edge, as can be seen in Fig. 5.D-E. The uncertainty of the cell center determination and the fact that two cells of a not perfectly round MB can have different r / R values even if they belong to the same concentric layer explain why the cell layer assignment is a more accurate determination of the cell location in the MB. ***: p < 0.001; N.S.: non significant.

**Supplementary Figure 4.**
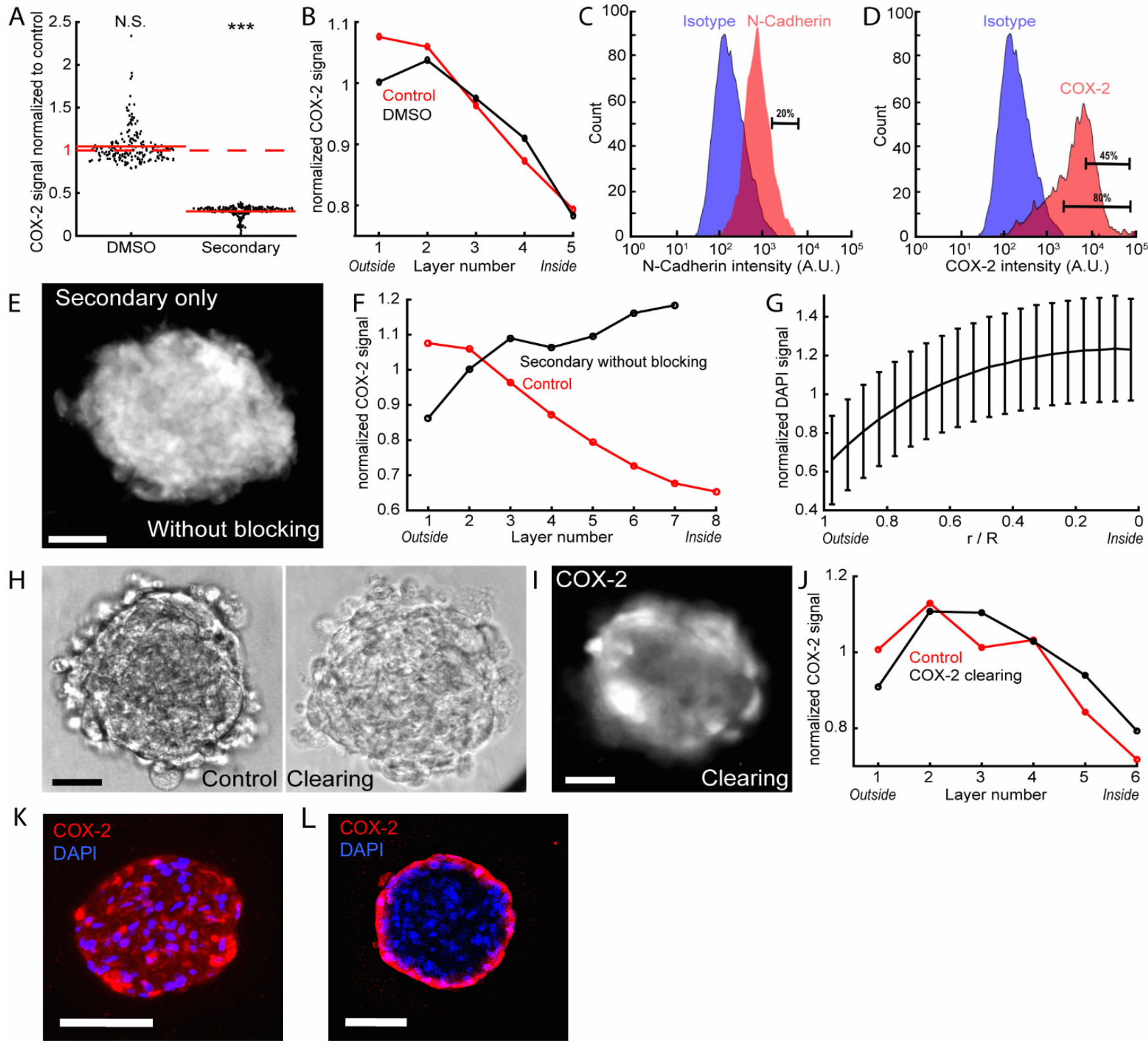
Validation of the fluorescent signal patterns. (A-B) Specificity of the staining and non-interference of the DMSO. Performing the immunostaining without the primary antibody and staining only with secondary antibody resulted in a very low fluorescent signal, which validated the specificity of the primary antibodies (A, control for secondary: N_chips_ = 1; n_MBs_ = 192; secondary: N_chips_ = 1; n_MBs_ = 251). MBs were formed and cultivated in the presence of 0.1 % (v/v) DMSO (the maximum concentration in culture media containing the inhibitors). Similar average per chip (A, control for DMSO: N_chips_ = 3; n_MBs_ = 644; DMSO 0.1 % (v/v): N_chips_ = 1; n_MBs_ = 195) and distribution of fluorescent signal in the cell layers (B, DMSO 0.1 % (v/v): n_cells_ = 5,982) demonstrated the absence of contribution of 0.1 % (v/v) DMSO in the cell behavior within MBs. (C-D) Flow cytometry analysis of the percentage of COX-2^high^ (C) and N-cadherin (D) expressing cells, after MBs dissociation and immunostaining, revealed the presence of several subpopulations expressing different levels of these proteins, however, without any spatial information (at least 5000 cells were analysis for each condition). (E-F) Removing the blocking step during the staining showed that there is no limitation for antibody diffusion. The cells were fixed, permeabilized and then stained only with the secondary antibody, without blocking the samples (E), rendering all immunogenic sites of the MBs accessible. Fluorescent signal distribution in the different cell layers demonstrated higher signal intensity in the core of MBs than in the edge (F, Control: N_chips_ = 13; n_MBs_ = 2,936; n_cells_ = 159,596; Secondary without blocking: N_chips_ = 1; n_MBs_ = 17; n_cells_ = 1,618). (G-J) Quantifying the DAPI signal (G) and clearing the samples (H-J) showed that there was no significant light path alteration in the 3D MBs. (G) The DAPI fluorescent signal distribution inside the MBs displayed a continuous signal increase from the edge (r/R = 1) to the core (r/R = 0; N_chips_ = 55; n_MBs_ = 10,072; n_cells_ = 699,836), which demonstrated that their in no diffusion limitation of small molecules and that the fluorescent light path is not attenuated by the MB opacity. (H-J) The MBs were subjected to ClearT2 treatment after the immunostaining for COX-2. Representative images (H) showed that the MBs were efficiently cleared post ClearT2, but the distribution of the fluorescent signal intensity was not affected as demonstrated by a representative MB (I) and the quantification of the distribution of the fluorescent signal after clearing in the different cell layers (J, control COX-2: N_chips_ = 1; n_MBs_ = 23; n_cells_ = 2,366; clearing COX-2: N_chips_ = 1; n_MBs_ = 67; n_cells_ = 6,333). The MBs were recovered from the chip then cryosectionned at 7 μm. For this cell layer depth, there is no antibody diffusion limitation or light path alteration. The COX-2 distribution signal show similar pattern as obtained with wide field imaging (K). Alternatively, the MBs were image using a 2-photons microscope and the COX-2 fluorescent signal pattern show similar distribution as wide field imaging (L). All scale bars are 50 μm. These results demonstrated the reliability of the measurements by image analysis, ensuring (1) the specificity of the fluorescent labeling; (2) the absence of limitation for antibody diffusion; (3) the absence of the light path alteration in the 3D structures.

**Supplementary Figure 5.**
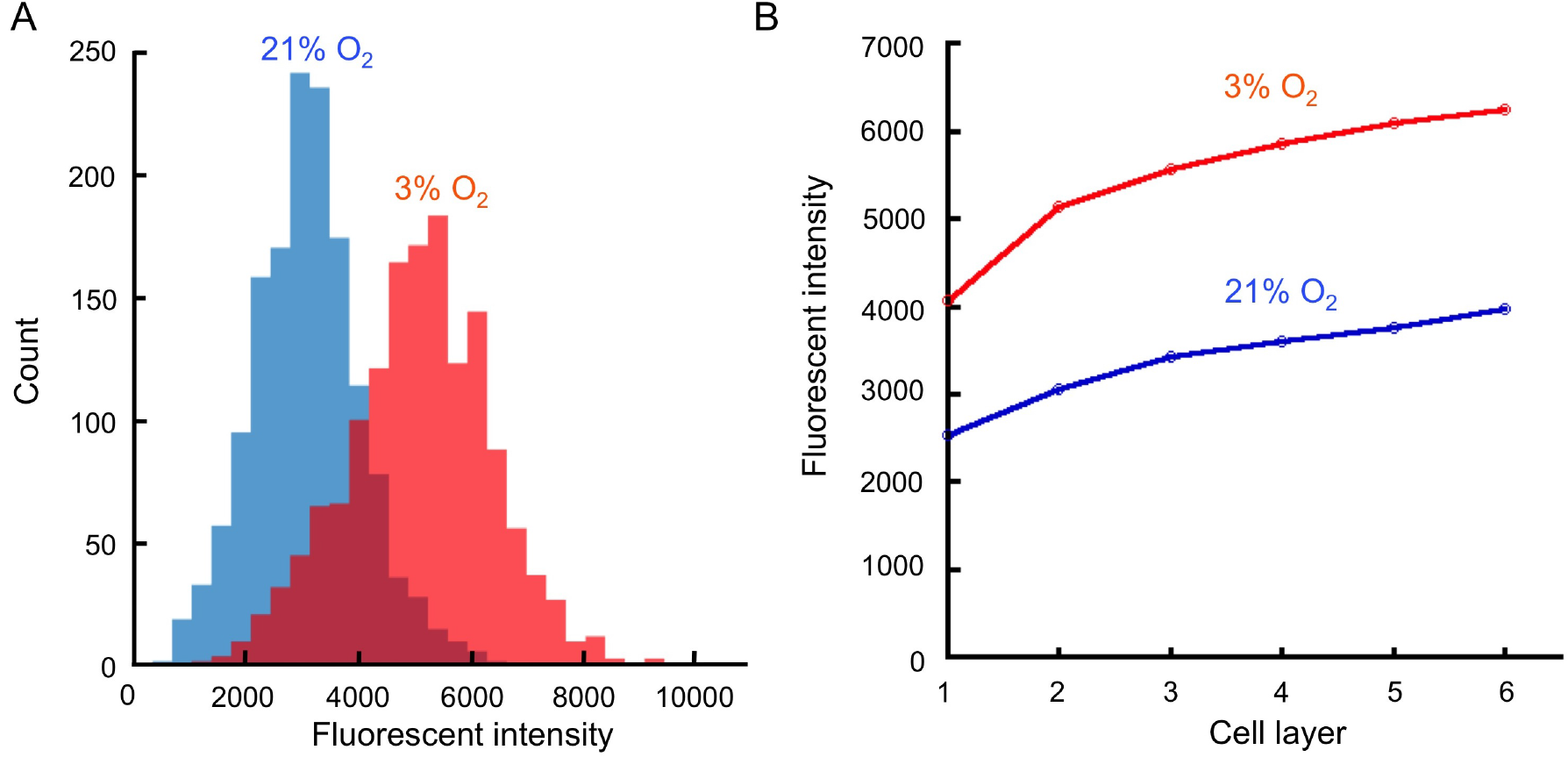
Hypoxia analysis within mesenchymal bodies. Mesenchymal bodies were incubated with Image-iT™ Red Hypoxia Reagent and placed in incubators set up at 21% or 3% O_2_. After 24h, a higher number of cells was positive for Image-iT™ Red Hypoxia Reagent at 3% than at 21% O_2_ (A). While a slight increase in hypoxia signal was detected in the inner cell layers at 21%O_2_, this fluorescent signal never reached the level measured at 3% O_2_, even at the most outer layers, demonstrating the absence of hypoxic core in the MBs.

**Supplementary Figure 6.**
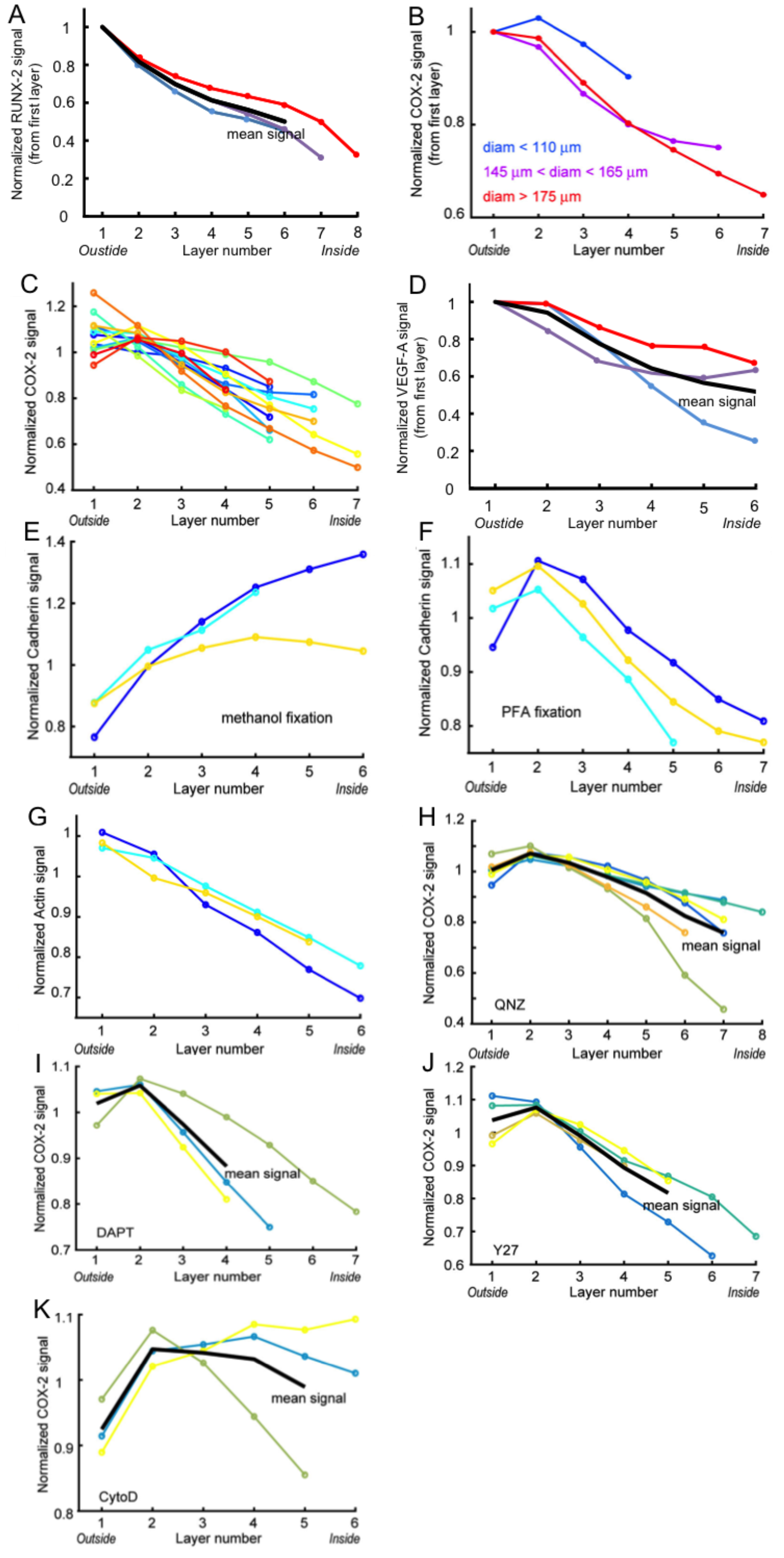
Intra-MB fluorescent signal distribution in individual chip. Evolution of the intra-MB fluorescent signal with control conditions (A-E, see Fig. 3 in the main text) and inhibitors (F-I, see Fig. 5 in the main text). (A) Normalized mean COX-2 with the cell layer number for MBs having different sizes (N_chips_ = 13; blue, diameter < 110 μm, n_MBs_ = 298, n_cells_ = 9,282; purple, 145 μm< diameter < 165 μm, n_MBs_ = 620, n_cells_ = 42,469; red, diameter > 175 μm, n_MBs_ = 295, n_cells_ = 26,251) in control conditions. (B-E) Normalized mean COX-2 (B N_chips_ = 13, n_MBs_ = 2,936, n_cells_ = 159,596), cadherin (C, with methanol fixation, N_chips_ = 3, n_MBs_ = 405, n_cells_ = 24,185; D, with PFA fixation, N_chips_ = 3, n_MBs_ = 649, n_cells_ = 47,254) and actin (E, N_chips_ = 3, n_MBs_ = 421, n_cells_ = 23,970) signals with the cell layer number. Each color represents the mean behavior for one chip. (F-I) Evolution of the normalized mean COX-2 signal with the cell layer number depending on the inhibitor: QNZ (F, N_chips_ = 6, n_MBs_ = 1,215, n_cells_ = 117,443), DAPT (G, N_chips_ = 3, n_MBs_ = 658, n_cells_ = 37,165), Y27 (H, N_chips_ = 4, n_MBs_ = 709, n_cells_ = 45,839) or CytoD (I, N_chips_ = 3, n_MBs_ = 458, n_cells_ = 28,981). Each color represents the mean behavior for one chip. The black lines represent the mean of the single chips.

**Supplementary Figure 7.**
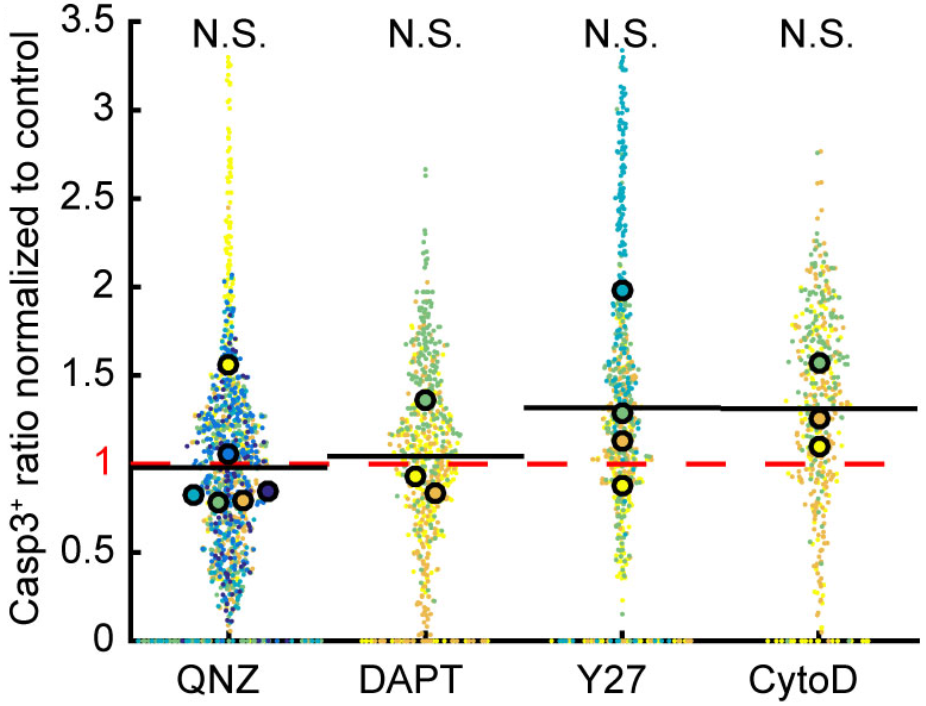
Ratio of Casp3^+^ cells per MBs formed with QNZ (n_MBs_ = 1,216), DAPT (n_MBs_ = 658), Y-27 (n_MBs_ = 709) and CytoD (n_MBs_ = 458). For each of these experiments, the results are normalized by the mean of the corresponding control. N.S.: not significant; **: p < 0.01.

## REFERENCES

1. Rossi, G., Manfrin, A. & Lutolf, M. P. Progress and potential in organoid research. Nat. Rev. Genet. 19, 671–687 (2018).

2. Singer, Z. S. et al. Dynamic Heterogeneity and DNA Methylation in Embryonic Stem Cells. Mol Cell 55, 319–331 (2014).

3. Hough, S. R. et al. Single-Cell Gene Expression Profiles Define Self-Renewing, Pluripotent, and Lineage Primed States of Human Pluripotent Stem Cells. Stem Cell Reports 2, 881–895 (2014).

4. Kim, T.-H. et al. Single-Cell Transcript Profiles Reveal Multilineage Priming in Early Progenitors Derived from Lgr5 + Intestinal Stem Cells. Cell Reports 16, 2053–2060 (2016).

5. Turner, D. A., Baillie-Johnson, P. & Martinez Arias, A. Organoids and the genetically encoded self-assembly of embryonic stem cells. Bioessays 38, 181–191 (2016).

6. Takebe, T., Zhang, B. & Radisic, M. Synergistic Engineering: Organoids Meet Organs-on-a-Chip. Cell Stem Cell 21, 297–300 (2017).

7. Dominici, M. et al. Minimal criteria for defining multipotent mesenchymal stromal cells. The International Society for Cellular Therapy position statement. Cytotherapy 8, 315–317 (2006).

8. Jeon, S. et al. Shift of EMT gradient in 3D spheroid MSCs for activation of mesenchymal niche function. Scientific Reports 7, 6859 (2017).

9. Freeman, B. T., Jung, J. P. & Ogle, B. M. Single-Cell RNA-Seq of Bone Marrow-Derived Mesenchymal Stem Cells Reveals Unique Profiles of Lineage Priming. PLoS ONE 10, e0136199 (2015).

10. Lee, W. C. et al. Multivariate biophysical markers predictive of mesenchymal stromal cell multipotency. PNAS 111, E4409–E4418 (2014).

11. Bartosh, T. J. et al. Aggregation of human mesenchymal stromal cells (MSCs) into 3D spheroids enhances their antiinflammatory properties. Proc. Natl. Acad. Sci. U.S.A. 107, 13724–13729 (2010).

12. Bhumiratana, S. et al. Large, stratified, and mechanically functional human cartilage grown in vitro by mesenchymal condensation. Proc. Natl. Acad. Sci. U.S.A. 111, 6940–6945 (2014).

13. Zelzer, E. & Olsen, B. R. The genetic basis for skeletal diseases. Nature 423, 343–348 (2003).

14. Zelzer, E. et al. VEGFA is necessary for chondrocyte survival during bone development. Development 131, 2161–2171 (2004).

15. Biddulph, D. M., Dozier, M. M. & Capehart, A. A. Inhibition of prostaglandin synthesis reduces cyclic AMP levels and inhibits chondrogenesis in cultured chick limb mesenchyme. Methods Cell Sci 22, 9–16 (2000).

16. Huang, C. et al. The spatiotemporal role of COX-2 in osteogenic and chondrogenic differentiation of periosteum-derived mesenchymal progenitors in fracture repair. PLoS ONE 9, e100079 (2014).

17. Rundle, C. H. et al. Retroviral-based gene therapy with cyclooxygenase-2 promotes the union of bony callus tissues and accelerates fracture healing in the rat. J Gene Med 10, 229–241 (2008).

18. Zhang, X. et al. Cyclooxygenase-2 regulates mesenchymal cell differentiation into the osteoblast lineage and is critically involved in bone repair. J. Clin. Invest. 109, 1405–1415 (2002).

19. Mammoto, T. et al. Mechanochemical control of mesenchymal condensation and embryonic tooth organ formation. Dev. Cell 21, 758–769 (2011).

20. Abbyad, P., Dangla, R., Alexandrou, A. & Baroud, C. N. Rails and anchors: guiding and trapping droplet microreactors in two dimensions. Lab Chip 11, 813–821 (2011).

21. Sart, S., Tomasi, R. F.-X., Amselem, G. & Baroud, C. N. Multiscale cytometry and regulation of 3D cell cultures on a chip. Nat Commun 8, 469 (2017).

22. Delorme, B. et al. Specific plasma membrane protein phenotype of culture-amplified and native human bone marrow mesenchymal stem cells. Blood 111, 2631–2635 (2008).

23. Jin, H. J. et al. Down-regulation of CD105 is associated with multi-lineage differentiation in human umbilical cord blood-derived mesenchymal stem cells. Biochemical and Biophysical Research Communications 381, 676–681 (2009).

24. Russell, K. C. et al. In vitro high-capacity assay to quantify the clonal heterogeneity in trilineage potential of mesenchymal stem cells reveals a complex hierarchy of lineage commitment. Stem Cells 28, 788–798 (2010).

25. Jones, M. et al. CD146 expression, as a surrogate biomarker for human mesenchymal stromal cell multilineage differentiation, is preserved during cell expansion in an automated hollow-fiber membrane bioreactor. PHARMACEUTICAL BIOPROCESSING 6, 93–105 (2018).

26. Sacchetti, B. et al. Self-renewing osteoprogenitors in bone marrow sinusoids can organize a hematopoietic microenvironment. Cell 131, 324–336 (2007).

27. Honda, H. Description of cellular patterns by Dirichlet domains: the two-dimensional case. J. Theor. Biol. 72, 523–543 (1978).

28. Chang, H., Yang, Q. & Parvin, B. Segmentation of heterogeneous blob objects through voting and level set formulation. Pattern Recognit Lett 28, 1781–1787 (2007).

29. Jin, H. J. et al. Downregulation of Melanoma Cell Adhesion Molecule (MCAM/CD146) Accelerates Cellular Senescence in Human Umbilical Cord Blood-Derived Mesenchymal Stem Cells. Stem Cells Transl Med 5, 427–439 (2016).

30. Alimperti, S. & Andreadis, S. T. CDH2 and CDH11 as Regulators of Stem Cell Fate Decisions. Stem Cell Res 14, 270–282 (2015).

31. Kawaguchi, J., Kii, I., Sugiyama, Y., Takeshita, S. & Kudo, A. The transition of cadherin expression in osteoblast differentiation from mesenchymal cells: consistent expression of cadherin-11 in osteoblast lineage. J. Bone Miner. Res. 16, 260–269 (2001).

32. Knudsen, K. A. & Soler, A. P. Cadherin-mediated cell-cell interactions. Methods Mol. Biol. 137, 409–440 (2000).

33. Monier-Gavelle, F. & Duband, J. L. Cross talk between adhesion molecules: control of N-cadherin activity by intracellular signals elicited by beta1 and beta3 integrins in migrating neural crest cells. J. Cell Biol. 137, 1663–1681 (1997).

34. Shore, E. M. & Nelson, W. J. Biosynthesis of the cell adhesion molecule uvomorulin (E-cadherin) in Madin-Darby canine kidney epithelial cells. J. Biol. Chem. 266, 19672–19680 (1991).

35. Menko, A. S., Zhang, L., Schiano, F., Kreidberg, J. A. & Kukuruzinska, M. A. Regulation of cadherin junctions during mouse submandibular gland development. Dev. Dyn. 224, 321–333 (2002).

36. McMillen, P. & Holley, S. A. Integration of cell-cell and cell-ECM adhesion in vertebrate morphogenesis. Curr. Opin. Cell Biol. 36, 48–53 (2015).

37. Cosgrove, B. D. et al. N-Cadherin adhesive interactions modulate matrix mechanosensing and fate commitment of mesenchymal stem cells. Nat Mater 15, 1297–1306 (2016).

38. Hinz, B., Pittet, P., Smith-Clerc, J., Chaponnier, C. & Meister, J.-J. Myofibroblast development is characterized by specific cell-cell adherens junctions. Mol. Biol. Cell 15, 4310–4320 (2004).

39. Boregowda, S. V., Booker, C. N. & Phinney, D. G. Mesenchymal Stem Cells: The Moniker Fits the Science. Stem Cells 36, 7–10 (2018).

40. Mahoney, D. J. et al. TSG-6 regulates bone remodeling through inhibition of osteoblastogenesis and osteoclast activation. J. Biol. Chem. 283, 25952–25962 (2008).

41. Yoshiko, Y., Aubin, J. E. & Maeda, N. Stanniocalcin 1 (STC1) protein and mRNA are developmentally regulated during embryonic mouse osteogenesis: the potential of stc1 as an autocrine/paracrine factor for osteoblast development and bone formation. J. Histochem. Cytochem. 50, 483–492 (2002).

42. Arikawa, T., Omura, K. & Morita, I. Regulation of bone morphogenetic protein-2 expression by endogenous prostaglandin E2 in human mesenchymal stem cells. J. Cell. Physiol. 200, 400–406 (2004).

43. Kuwano, T. et al. Cyclooxygenase 2 is a key enzyme for inflammatory cytokine-induced angiogenesis. FASEB J. 18, 300–310 (2004).

44. Zhu, Y. M., Azahri, N. S. M., Yu, D. C. W. & Woll, P. J. Effects of COX-2 inhibition on expression of vascular endothelial growth factor and interleukin-8 in lung cancer cells. BMC Cancer 8, 218 (2008).

45. Németh, K. et al. Bone marrow stromal cells attenuate sepsis via prostaglandin E(2)-dependent reprogramming of host macrophages to increase their interleukin-10 production. Nat. Med. 15, 42–49 (2009).

46. Gallardo-Pérez, J. C. et al. NF-kappa B is required for the development of tumor spheroids. J. Cell. Biochem. 108, 169–180 (2009).

47. Kipp, A. P., Deubel, S., Arnér, E. S. J. & Johansson, K. Time- and cell-resolved dynamics of redox-sensitive Nrf2, HIF and NF-κB activities in 3D spheroids enriched for cancer stem cells. Redox Biol 12, 403–409 (2017).

48. Susperregui, A. R. G. et al. Noncanonical BMP signaling regulates cyclooxygenase-2 transcription. Mol. Endocrinol. 25, 1006–1017 (2011).

49. Mehrotra, M., Saegusa, M., Voznesensky, O. & Pilbeam, C. Role of Cbfa1/Runx2 in the fluid shear stress induction of COX-2 in osteoblasts. Biochem. Biophys. Res. Commun. 341, 1225–1230 (2006).

50. Sero, J. E. et al. Cell shape and the microenvironment regulate nuclear translocation of NF-κB in breast epithelial and tumor cells. Mol. Syst. Biol. 11, 790 (2015).

51. Lee, E. J. et al. N-cadherin determines individual variations in the therapeutic efficacy of human umbilical cord blood-derived mesenchymal stem cells in a rat model of myocardial infarction. Mol. Ther. 20, 155–167 (2012).

52. Zhou, Y., Chen, H., Li, H. & Wu, Y. 3D culture increases pluripotent gene expression in mesenchymal stem cells through relaxation of cytoskeleton tension. J. Cell. Mol. Med. 21, 1073–1084 (2017).

53. Lana-Elola, E., Rice, R., Grigoriadis, A. E. & Rice, D. P. C. Cell fate specification during calvarial bone and suture development. Dev. Biol. 311, 335–346 (2007).

54. Boutin, M. E. & Hoffman-Kim, D. Application and assessment of optical clearing methods for imaging of tissue-engineered neural stem cell spheres. Tissue Eng Part C Methods 21, 292–302 (2015).

